# Identification of functional residues using machine learning provides insights into the evolution of odorant receptor gene families in solitary and social insects

**DOI:** 10.1101/2021.12.10.472060

**Authors:** Pablo Mier, Jean-Fred Fontaine, Marah Stoldt, Romain Libbrecht, Carlotta Martelli, Susanne Foitzik, Miguel A. Andrade-Navarro

**Author notes:** Corresponding Author: Pablo Mier.

## Abstract

The gene family of insect odorant receptors (ORs) has greatly expanded in the course of evolution. ORs allow insects to detect volatile chemicals and therefore play an important role in social interactions, the detection of enemies and preys, and during foraging. The sequences of several thousand ORs are known, but their specific function or ligands have been identified only for very few of them. To advance the functional characterization of ORs, we compiled, curated and aligned the sequences of 3,902 ORs from 21 insect species. We identified the amino acid positions that best predict the response to ligands using machine learning on sets of functionally characterized proteins from the fly *Drosophila melanogaster*, the mosquito *Anopheles gambiae* and the ant *Harpegnathos saltator*. We studied the conservation of these predicted relevant residues across all OR subfamilies and show that the subfamilies that expanded strongly in social insects exhibit high levels of conservation in their binding sites. This indicates that ORs of social insect families are typically finely tuned and exhibit a sensitivity to very similar odorants. Our novel approach provides a powerful tool to use functional information from a limited number of genes to investigate the functional evolution of large gene families.

## Introduction

Odorant receptors (ORs) constitute the largest family of chemoreceptors expressed in the membranes of olfactory receptor neurons in insects. The insect odorant receptor gene family is an evolutionary novelty in the ancestor of all insects [1], likely an adaptation to sensory perception in terrestrial life. Insects use ORs to perceive sexual pheromones, food sources including nectar-providing flowers and, importantly, for social communication [2-5].

A rapid expansion of chemoreceptors, in particular of ORs, accompanied in Hymenoptera the repeated transition from a solitary to a social lifestyle [6,7]. The ecological success of social insects is based on their ability to form complex cooperative societies, which in turn was only made possible by their sophisticated chemical communication [8-10]. Especially fascinating and diverse communication mechanisms can be found in the ants. Ants use secretions of 70 different glands to exchange information with their nestmates and in addition signal their colony membership, fecundity and caste via a complex mix of long-chained hydrocarbons on their cuticle (CHCs) [11]. As in all other insects, the antenna is the primary organ for odorant perception, where ants can express up to 500 different ORs in olfactory sensory neurons [12]. The connection between sociality and odorant receptor repertoire in ants is furthermore supported by experimental studies showing that the impairment of certain receptors affects social behavior [13,14], and by the finding that the partial loss of social behaviors in social parasites was accompanied by a loss of OR genes [15,16]. The 9-exon subfamily shows a particularly strong signal of expansion and association with the switch to sociality and social communication and this not only in the ants [17-21], but also in the social wasps [5]. As some 9-exon ORs bind multiple ligands and some bind the same [22,23], it has been suggested that this subfamily distinguishes odors using combinatorial coding [5,24]. Together with the aforementioned expansion of this subfamily this might allow some insects, including ants, to discriminate between a large variety of odors, probably CHCs. Thus, especially this OR subfamily is a prime candidate for understanding how advances in chemical communication led to the establishment of eusocial societies in ants. While the specificity and tuning of the different ORs has been well studied in dipteran model species such as *Drosophila melanogaster* and *Anopheles gambiae* [25-27], it remains largely unclear to which chemicals the large number of ORs in social insects respond to [23]. This knowledge would be necessary to make predictions about the trajectory leading to the evolution of eusociality in insects.

Here we present an approach to expand the functional characterization of the OR protein family by using machine learning. Machine learning has been used already in the field of insect ORs to identify ligands for specific ORs (e.g. [28,29]). Here, we use machine learning with the goal of providing functional and evolutionary hypotheses for the entire OR family. For this, we collected and curated the sequences of 3,902 OR proteins from 21 insect species. We constructed a multiple sequence alignment of these sequences. Then we used machine learning to evaluate the power of particular amino acid positions in the alignment to predict responses to chemicals according to available experimental data from three well-studied insects, the dipterans *Anopheles gambiae* [26] and *Drosophila melanogaster* [27], and the ant *H. saltator* [23]. Predictive amino acids were then mapped to 3D locations using as template the only solved structure of a protein from the OR family [30]. Independently from the machine learning approach, we used sequence similarity to cluster the OR families from 21 species into groups that are expected to have similar biological functions across different species. We annotated these clusters according to their evolutionary expansion, taxonomic specificity, and conservation of their predicted binding sites, to find modes of evolution associated to the emergence of biological and molecular function. Finally, as a first example on how our approach can be used to test biological hypotheses related to the evolution of eusociality, we tested whether gene subfamilies, which expanded in particular social insects are characterized by specific tuning patterns.

Our approach provides an easy way to transfer information between thousands of ORs already considered, and allows expanding this information either to individual ORs from genomes not yet included in our resource, or potentially by including relevant OR datasets from complete genomes as needed, as well as new functional profiles. This approach can be potentially applied to other large families of paralogs. Analysis of these large families should allow us to understand how events of gene duplication drive the emergence of novel functions.

## Materials and Methods

### Sequence data retrieval, curation and alignment

We obtained the annotated odorant receptor (OR) protein sequences from the following 21 insect species, including 8 ant species, 2 social bee species and 11 solitary insects from damsel flies to flies (**Table 1**): *Drosophila melanogaster* [31], *Anopheles gambiae* [26], *Apis mellifera, Solenopsis invicta, Nasonia vitripennis* and *Ooceraea biroi* [12], *Pogonomyrmex barbatus* [32], *Atta cephalotes* and *Acromyrmex echinatior* [18], *Camponotus floridanus* and *Harpegnathos saltator* [17], *Linepithema humile* [6], *Blattella germanica* [33], *Calopteryx splendens* [34], *Bombus terrestris* [35], *Tribolium castaneum* [36], *Cloeon dipterum* [37], *Manduca sexta* [38], *Pediculus humanus* [39], *Acyrthosiphon pisum* and *Aphis glycines* [40].

**Table 1.**
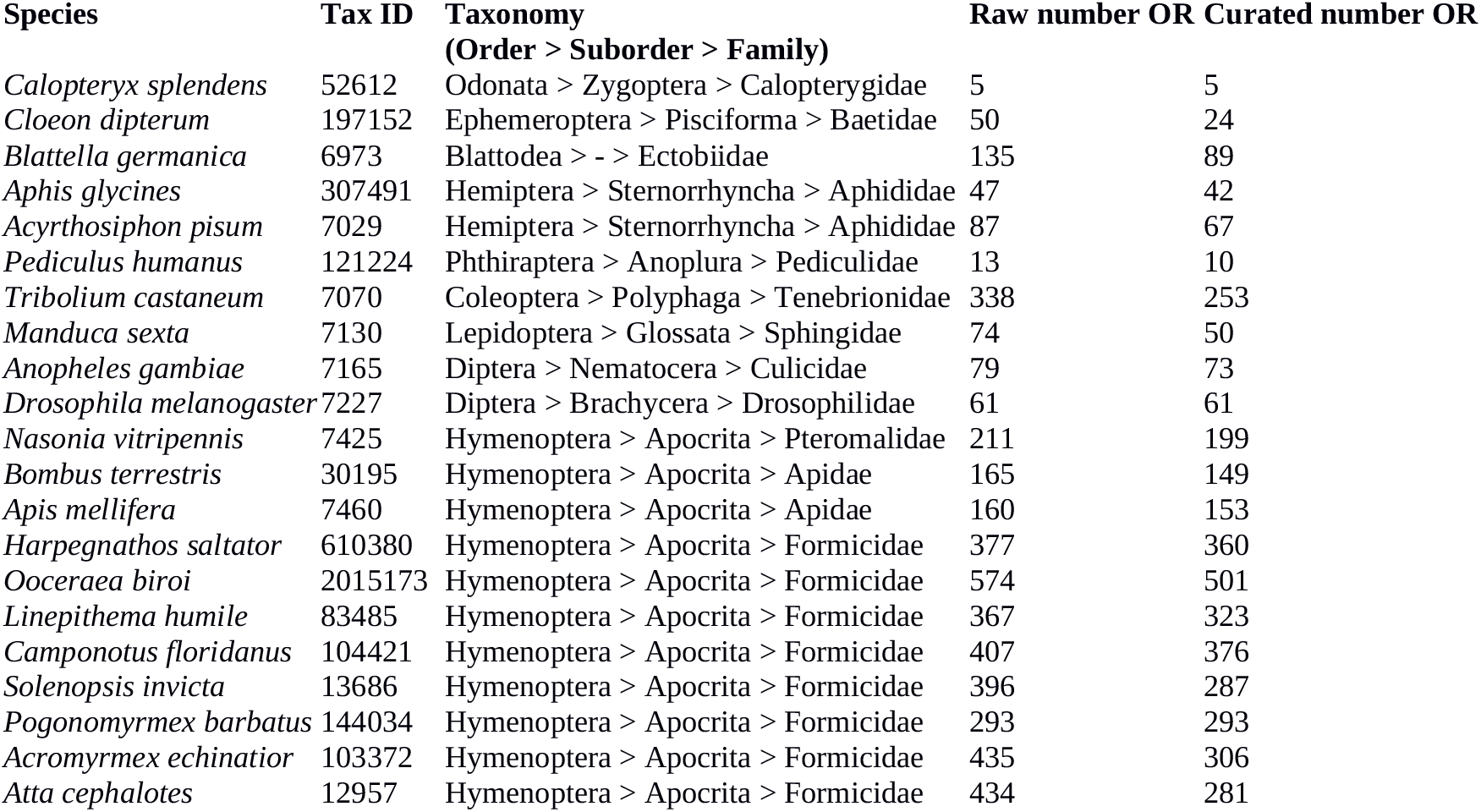
List of insect species used and number of raw and curated Odorant Receptors.

For the manual curation process, sequences from the retrieved raw datasets identified as pseudogenes or fragments were removed. After that, sequences showing large and unique insertions and deletions, or large stretches with low conservation compared to the aligned sequences were additionally removed. The full dataset of 3,902 curated OR proteins was aligned using MAFFT v7.453 with default parameters [41] (**Supplementary File S1**). The complete taxonomic lineage from each of the species was obtained from the NCBI resource Common Taxonomy Tree [42]. We resolved the phylogenetic relationships in ants with information from [43].

### Clustering of OR proteins

Protein sets were clustered using a method designed to find groups of proteins with similar functions across species (OrthoFinder v2.3.12 with default parameters [44]). These clusters were compared to the subfamilies in *C. floridanus* and *H. saltator* provided by [17], and the ones from *A. echinatior* and *A. cephalotes* provided by [18]. In *A. cephalotes*, we renamed the subfamily “unassigned N???” into “unassigned”, to be consistent with the unassigned records in the other species. Similarly, missing subfamily information was denoted as “unassigned”. Additionally, in *A. echinatior* one OR was annotated with subfamily “0” (typo in the original publication), and we changed it to “O”.

### Machine learning approach

The machine learning procedure uses as input a table created using the multiple sequence alignment of all the OR sequences. In this table, amino acids (table’s cells) of proteins (rows) can be seen at each position of the alignment (columns, or machine learning variables). A machine learning variable is defined here as a position in the alignment (column in the multiple sequence alignment) and a machine learning feature as an amino acid at a specific position. Additional columns include numerical chemical-response data for some proteins from datasets from three species: *D. melanogaster* [27], *A. gambiae* [26], and *H. saltator* [23]. For the machine learning training phase, 8 out of 672 chemicals with the highest numbers of values (>100 values in the union of the 3 datasets) and 494 out of 3,902 proteins (associated with at least one chemical-response value) were selected. The eight selected chemicals have the following registry numbers and IUPAC names (common names indicated within parentheses): 108-94-1 cyclohexanone, 431-03-8 butane-2,3-dione (diacetyl), 67-64-1 propan-2-one (acetone), 110-43-0 heptan-2-one (2-heptanone), 6728-26-3 (E)-hex-2-enal (trans-2-hexenal), 119-36-8 methyl 2-hydroxybenzoate (methyl salicylate), 105-87-3 [(2E)-3,7-dimethylocta-2,6-dienyl] acetate (geranyl acetate), 3391-86-4 oct-1-en-3-ol (vinyl amyl carbinol).

For each chemical within each dataset, chemical-response values were binarized by setting a value greater than the 75th percentile to one to represent positive response, 0 otherwise to represent lack of response (**Supplementary File S2;** not-tested combinations simply lack values). For each chemical-dataset pair (3 datasets and 8 chemicals: 24 pairs), a random forest (RF) model based on 500 trees was trained to predict chemical-response values using the machine learning variables. Only proteins associated with a chemical-response value were used in the training set. Also, near zero-variance variables were filtered out. The analysis was implemented in R with the caret and randomForest packages (the optimal mtry parameter, defining the optimal number of predictors for split, was defined by grid search during training phase; tested mtry values: 20, 50 and 100). A model’s performance was derived from internal cross-validations (10-fold cross-validations repeated 10 times) and model’s measures of feature importance were scaled by the caret package to have a maximum value of 100. Performance during the cross-validations is reported as area under ROC curves, F1 score, sensitivity or precision.

### Computation of sequence conservation

To measure the sequence conservation of ORs in a cluster, each position in the alignment received a degree of conservation, which is simply the occurrence of the most frequent residue at the position for the ORs in the cluster: a conservation value of one indicates a totally conserved position, while very variable positions receive values close to zero. Then, for each cluster, we calculated a background sequence conservation, which is the average conservation value of all the residues of the sequence, and for comparison, a predictive residue conservation, which is the average conservation of predictive residues selected by machine learning. In general, we restricted this computation to clusters with five or more ORs.

## Results and Discussion

### Collection and curation of insect Odorant Receptor proteins

We started out by collecting the sequences of OR proteins from a diverse set of insect species. Furthermore, the OR protein sequences themselves are often mis-annotated, contain fragments or not translated sequences, or are described but not publicly available. To address these problems and build a robust and thorough dataset of insect OR proteins, we manually examined published data for 21 insect species with completely sequenced genomes (see Methods). Given the dynamic nature of the sequencing of new genomes, it seems necessary that such a collection could be updated as it would be of interest not only to other researchers in the field of OR evolution, but also to computational biologists developing methods for functional prediction using machine learning or other approaches. For these reasons, we developed a dedicated repository called iOrME (insect Odorant Receptors Molecular Evolution) to gather all raw and curated OR datasets, as well as taxonomical information for the insect species we used. It is available at http://cbdm-01.zdv.unimainz.de/~munoz/iorme/, with no restrictions for users.

For this first version of iOrME (v1.0) we collected a raw dataset of 4,708 OR sequences. The dataset also contained fragments, pseudogenes and sequences with sequencing errors. After manual curation, we ended up with a core dataset of 3,902 OR proteins (see Methods for details; **Figure 1; Table 1**). Some sets needed more curation than others, depending on the original work they were published in or their general scientific interest. For example, while for the leafcutter ant *Atta cephalotes* we removed 35% of the original sequences (from 434 to 281 proteins) and 52% for the mayfly *Cloeon dipterum* (from 50 to 24 proteins), the 61 well-established OR proteins from *D. melanogaster* remained, as well as the 293 proteins from the red harvester ant *Pogonomyrmex barbatus*.

**Figure 1.**
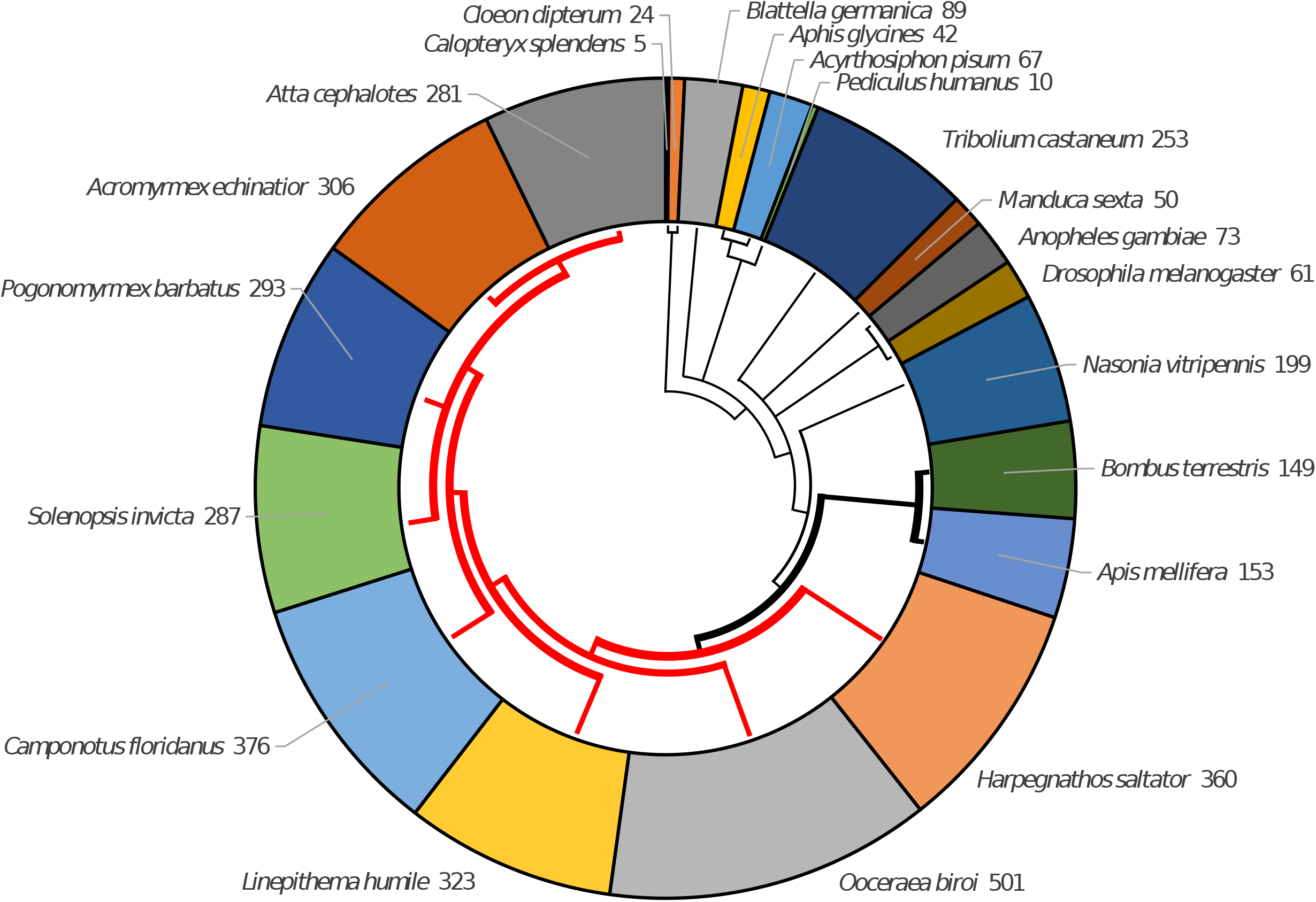
Number of ORs for each species. The tree represents the phylogenetic relationships between the species. Thick branches indicate social insects and red color indicates ants. While all social insects have more than 100 ORs, it is the case of only 2 non-social insects out of 11 (the wasp *N. vitripennis* and the beetle *T. castaneum*).

### The taxonomic distribution of ORs in clusters shows taxa-specific expansions

We performed sequence clustering of the 3,902 OR sequences, using a method that tries to group together proteins likely to perform similar functions across species (see Methods). Our goal is to evaluate the relation between the evolutionary history of these OR subfamilies and their ligand binding properties. The clustering yielded a total of 206 clusters, 40 of which consisted of a single protein (singletons) (**Supplementary File S3; Supplementary File S4;** FASTA files with the sequences of each cluster are available for download in iOrME). The largest clusters mostly correspond to subfamilies of ant ORs previously defined in [17] based on genome organization, and then expanded in [18] (**Supplementary Figure S1**).

Next, we evaluated the species distribution of the ORs (**Figure 2**). The taxonomic distribution of ORs is very variable across clusters reflecting the intricate evolutionary history of this family. Only two groups, C0 and C6 (with 589 and 138 sequences, respectively), have at least one protein from all of the 21 species. C6 includes Orco, one of the ancestral proteins of the family, very conserved across species, which forms a heteromeric cation channel with an OR subunit [45,46].

**Figure 2.**
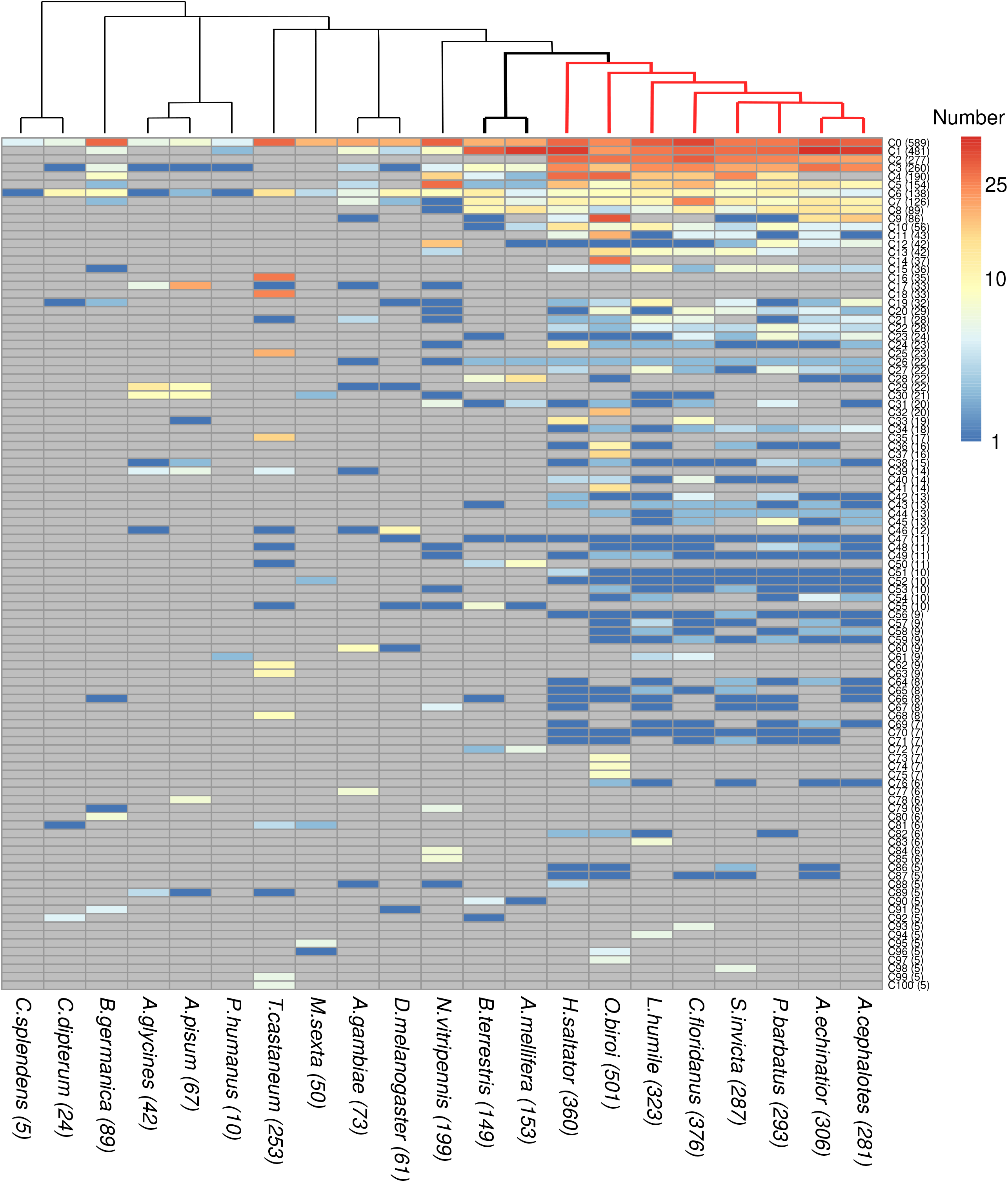
Species distribution of OR proteins per cluster. For each cluster, the number of ORs from each species is indicated. Only clusters with five or more proteins are shown. The tree above shows the phylogenetic relations of the 21 insect species: bold and red branches indicate social insects and ants, respectively. The total number of ORs per cluster and per species are shown in parenthesis.

Interestingly, one of the most populated clusters, C2 with 277 sequences, has ORs from all ants and only from the ants. It is comprised mostly of 9-exon ORs (**Supplementary Figure S1**), a subfamily known to be expanded in ants and paper wasps (not included in our dataset) [5,17-19,21]. OR genes in the 9-exon family were convergently lost in socially parasitic slavemaking ants [15]. Cluster C22 is ant specific too, in this case composed only of ORs from the V subfamily, also shown to be expanded in ants [18]. Among the single species clusters, C14 (37 sequences) and C16 (35 sequences), from the ant *O. biroi* and the beetle *T. castaneum*, respectively, stand out.

Analysis of these profiles can be used to investigate gene loss when a cluster has members from all but one or few species from one taxon. One such example is C40, which contains 14 OR genes from six of the eight ant species considered in our study but missing from *A. echinatior* and *A. cephalotes*, suggesting that this OR cluster was lost in the fungus-growing ants (Attini). Some clusters contain ORs from widely different species, which however did not expand. An extreme example is C47, which has 11 sequences from 11 species (*D. melanogaster* and the 10 Aculeata considered in this work, which includes the ant and bee species).

To evaluate the existence of taxa-specific expansions within our clusters, we measured the enrichment of taxon-specific ORs in each cluster, by computing for each cluster and taxon the log2-transformed ratio between the number of sequences from the given taxon and the number of species in it (**Figure 3**; for definitions see **Table 2**). Using this representation, we can find a number of clusters that reflect taxa-specific expansions in Hemiptera coupled to gene loss in ants: C17, C29 and C30. Many of these sequences were noted in [47] as the “Clade A” of *A. pisum*-specific recent and rapid OR expansion. We note also C28 (22 sequences) as the cluster with the most relevant Apoidea-specific expansion (13 sequences from *A. mellifera* and 6 from *B. terrestris*).

**Figure 3.**
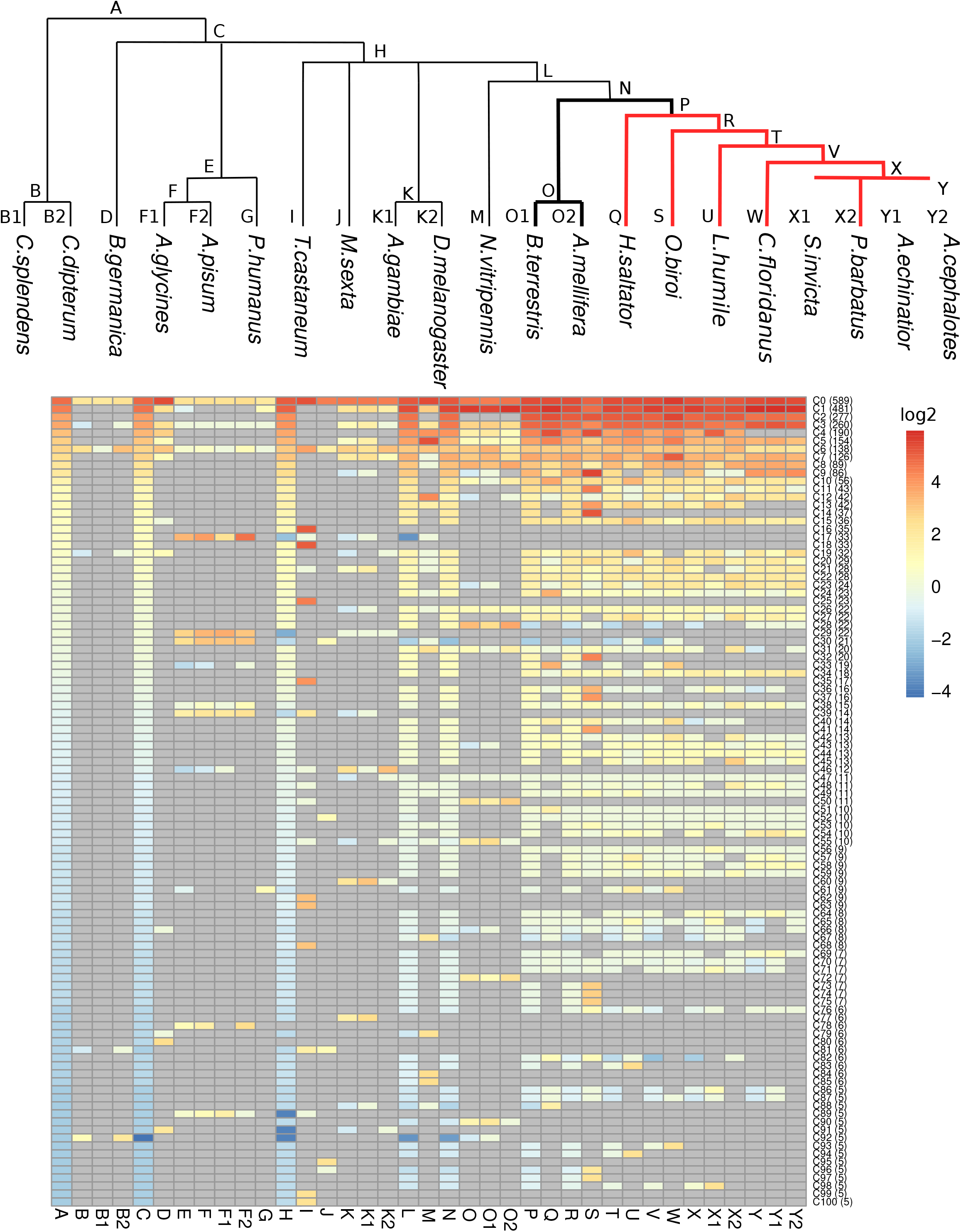
Enrichment of taxon-related ORs per cluster. For each cluster and each taxon, the log2-transformed ratio between the number of taxon-related OR proteins and the number of species in the taxon is shown. A positive value denotes a higher number of taxon-related proteins from the cluster than the number of species in the taxon. A negative value denotes a lower number of taxon-related proteins from the cluster than the number of species in the taxon. Each taxon is defined by a letter, depicted in the phylogenetic tree above and described in **Table 2**. In bold, social insects. In red, ants. Only clusters with five or more proteins are shown.

**Table 2.** Keys for taxonomic labels.

| **Code** | **Taxon** |
| --- | --- |
| [A] | Insecta |
| [B] | Palaeoptera |
| [B1] | *C. splendens* |
| [B2] | *C. dipterum* |
| [C] | Neoptera |
| [D] | *B. germanica* |
| [E] | Paraneoptera |
| [F] | Hemiptera |
| [F1] | *A. glycines* |
| [F2] | *A. pisum* |
| [G] | *P. humanus* |
| [H] | Endopterygota |
| [I] | *T. castaneum* |
| [J] | *M. sexta* |
| [K] | Diptera |
| [K1] | *A. gambiae* |
| [K2] | *D. melanogaster* |
| [L] | Hymenoptera |
| [M] | *N. vitripennis* |
| [N] | Aculeata |
| [O] | Apoidea |
| [O1] | *B. terrestris* |
| [O2] | *A. mellifera* |
| [P] | Formicoidea |
| [Q] | *H. saltator* |
| [R] | Formicoids |
| [S] | *O. biroi* |
| [T] | Formicoids - *O. biroi* |
| [U] | *L. humile* |
| [V] | Myrmicinae + *C. floridanus* |
| [W] | *C. floridanus* |
| [X] | Myrmicinae |
| [X1] | *S. invicta* |
| [X2] | *P. barbatus* |
| [Y] | Attini |
| [Y1] | *A. echinator* |
| [Y2] | *A. cephalotes* |

### Prediction of OR amino acid residues important for chemical binding following a machine learning approach

Insect odorant receptors bind chemicals to trigger neuronal activity essential for odorant perception [12]. While the functional information regarding chemical binding of the OR family is very limited, the multiple sequence alignment of the family contains a wealth of information about the residue variability in positions that include those interacting with the odorants. We hypothesized that given the availability of a few datasets profiling the neural response triggered by relatively large numbers of ORs upon standard sets of chemicals, using those with a machine learning approach would allow us to identify positions in the alignment corresponding to residues involved in the molecular recognition of odorants. Such an approach is supported by work that proposes that the entire OR family has a common mechanism of interacting with odorants according to structural analysis [48]. We used previously published data of the OR response to panels of chemicals from three insect species *D. melanogaster* (48 ORs, 618 chemicals) [27], *A. gambiae* (50 ORs, 110 chemicals) [26] and *H. saltator* (25 ORs, 37 chemicals) [23]. To identify and characterize amino acid positions and residues potentially important for the binding, we used both the available chemical binding information of eight selected chemicals (**Figure 4A**) and the OR sequence alignment to train machine learning models of prediction (see Methods for details). Classification performance varied across models during cross-validations, with often higher sensitivity than precision (**Figure 4B; Supplementary Figure S2; Supplementary File S5**). The predictions for some chemicals (e.g., 2,3-butanedione) are clearly better than for others. We also observe differences between datasets with generally worse predictions for the ant dataset: these can be due to the choice of chemicals.

**Figure 4.**
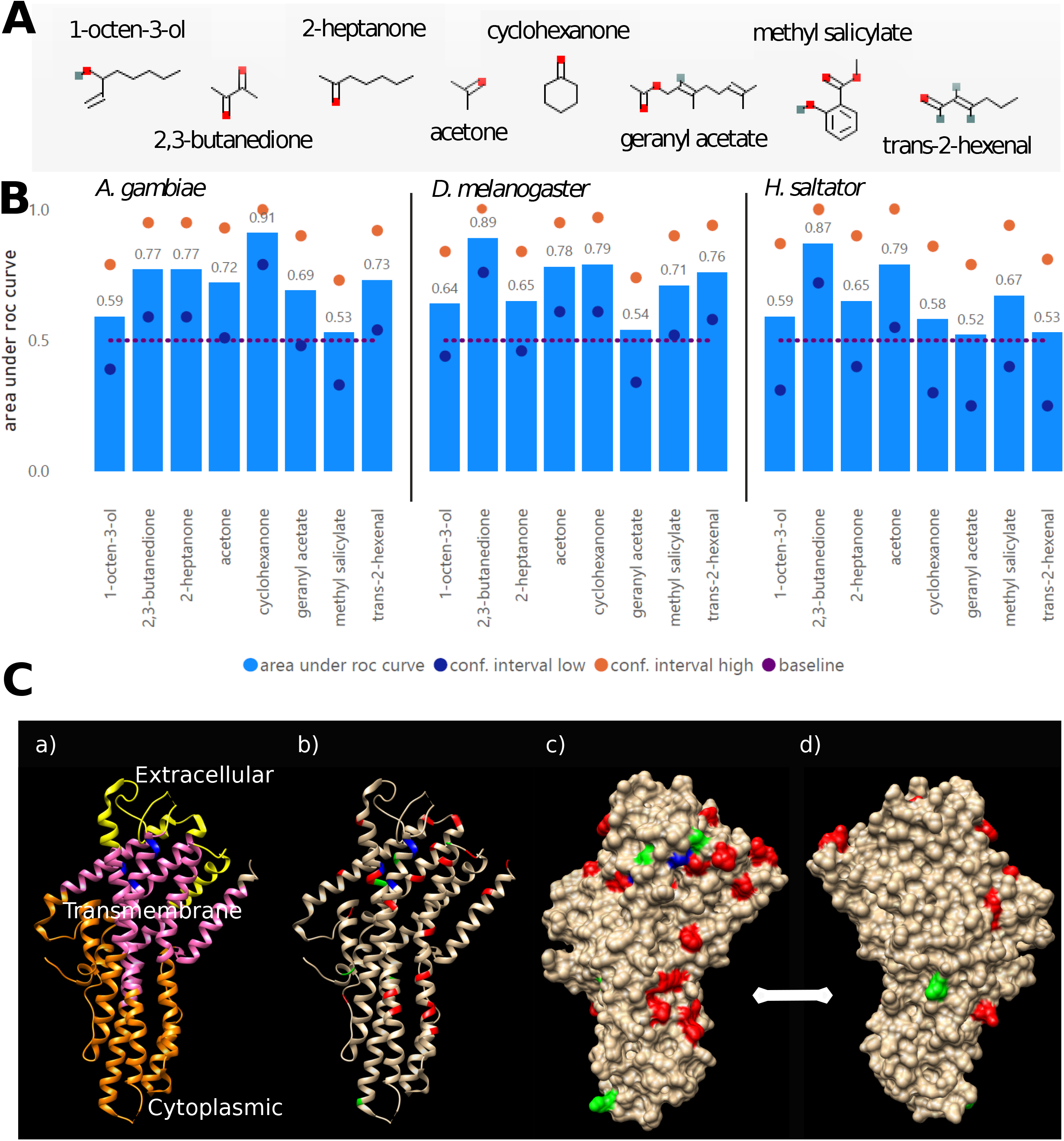
Results from the machine learning approach. **(A)** Chemicals used in the training (see Methods for registry names and IUPAC names). **(B)** Cross-validation performance. Performance of the machine learning models was evaluated by area under ROC curves (true positive rate against false positive rate at variable thresholds; best curve evaluating either the positive or the negative class) during 10-fold cross-validations repeated 10 times. A random forest model of 500 trees was trained for each species-chemical pair to predict the binding of the chemical to the species-related OR proteins. Low and high boundaries of the 95% confidence interval and baseline (0.5 = random classification) are shown. **(C)** Mapping predictive features on 3D structure: a) 3D structure of the Orco protein from the parasitic fig wasp *Apocrypta bakeri* (PDB:6C70) [33]; b)-d) Top 10 positions predicted for any of the three datasets are indicated in red. Positions detected among the top 100 in the three datasets (6 positions) are indicated in green. Positions indicated in blue (*A. bakeri* amino acid positions 143, 149-150, 202) were mapped from positions whose mutations were experimentally shown to modify ligand detection [49-51]; d) shows the molecule rotated 180° along the vertical axis.

After analyzing 2,892 distinct positions in the alignment, we obtained the importance of each amino acid at given positions (values range from 0 to 100), for each of the three datasets and for each of the eight chemicals (**Supplementary File S6**). Note that multiple amino acids can be found as predictive features for the same amino acid position, chemical and dataset. For example, T, P and K at position 1472 were found to be predictive for response to methyl salicylate, 119-36-8, for the *D. melanogaster* model, with importance of 31.49, 30.12 and 14.41, respectively, whereas at the same position and dataset, L, F and A were predictive for response to geranyl acetate, 105-87-3, with importance of 41.01, 27.32 and 26.79, respectively.

We observed that some positions were more often identified as predictive than others and we took this as evidence of their involvement in the molecular function of the OR family in general. **Table 3** lists the 10 most frequently found in each dataset. These were selected among those with importance > 10 and AUC > 0.7; a total of 475, 457, and 151, for the datasets from *D. melanogaster, A. gambiae* and *H. saltator*, respectively). As it could be expected, the more predictive features we observed, the more their positions were repeated and the better the performance of the machine learning method.

**Table 3.** Top predictive positions in the multiple sequence alignment as obtained from the machine learning approach per dataset. We mapped the top predictive positions from the multiple sequence alignment to the sequence of the ORCO_DROME protein (UniProtKB:Q9VNB5), and to its homologous protein B0FAQ4_APOBA (UniProtKB:B0FAQ4), for which there is an available 3D structure (PDB:6C70). Datasets: 1 = *D. melanogaster*, 2 = *A. gambiae*, 3 = *H. saltator*.

| Dataset | Times predictive | Alignment position | *D. melanogaster* Orco position | *A. bakeri* Orco position | Amino acid (D.m. / A.b.) |
| --- | --- | --- | --- | --- | --- |
| 1 | 8 | 1472 | 207 | 203 | L / V |
| 1 | 7 | 508 | 67 | 63 | N / E |
| 1 | 7 | 430 | 48 | 44 | V / V |
| 1 | 7 | 2529 | 414 | 402 | R / R |
| 1 | 7 | 2493 | 406 | 394 | F / F |
| 1 | 7 | 1069 | 143 | 139 | T / T |
| 1 | 7 | 2581 | 420 | 408 | S / S |
| 1 | 7 | 2855 | 486 | 474 | K / K |
| 1 | 7 | 620 | 83 | 79 | F / F |
| 1 | 7 | 1398 | 197 | 193 | I / F |
| 2 | 14 | 1210 | 170 | 166 | S / E |
| 2 | 11 | 1208 | 168 | 164 | T / T |
| 2 | 10 | 2345 | 387 | 375 | V / V |
| 2 | 10 | 430 | 48 | 44 | V / V |
| 2 | 9 | 1024 | - | - | - / - |
| 2 | 9 | 550 | 70 | 66 | E / D |
| 2 | 9 | 2591 | 421 | 409 | S / S |
| 2 | 9 | 2391 | 392 | 380 | F / A |
| 2 | 9 | 1594 | 229 | 225 | E / E |
| 2 | 9 | 334 | 30 | 25 | F / F |
| 3 | 2 | 216 | 16 | 11 | D / D |
| 3 | 2 | 1380 | - | - | - / - |
| 3 | 2 | 865 | 110 | 106 | Q / N |
| 3 | 2 | 1607 | 232 | 228 | Q / Q |
| 3 | 2 | 2771 | - | - | - / - |
| 3 | 2 | 2600 | 424 | 412 | E / E |
| 3 | 2 | 601 | 77 | 73 | N / N |
| 3 | 2 | 993 | - | - | - / - |
| 3 | 2 | 1474 | 208 | 204 | F / I |
| 3 | 2 | 2827 | 479 | 467 | F / F |

The top 10 predictive positions were mapped to the only available 3D structure for an insect OR (**Figure 4C**; PDB:6C70) [30], the Orco protein from the parasitic fig wasp *Apocrypta bakeri* (UniProtKB:B0FAQ4), using the sequence of the Orco protein from *D. melanogaster* (UniProtKB:Q9VNB5) as a link between the alignment of all ORs and the 3D structure. The ion channel structure is a heteromer of a specific OR with the OR co-receptor Orco [45,46], which opens upon ligand binding. Examination of the contact surface in the Orco tetrameric structure from *A. bakeri* suggests that the contact interface between subunits is in the lower-right part of the protein as displayed in **Figure 4C**. While a patch of positions overlaps the region of subunit interaction, most of the positions are in the top domain surrounding positions known to be close to the ligand binding pocket (in blue in **Figure 4C**; mapped from [49-51]).

A representation of the amino acid used for each of the positions in the clusters is provided as **Supplementary File S7**. The correspondence between the position in the alignment and those in Orco from *A. bakeri* is indicated in **Table 3** (all positions mapped in **Supplementary File S8**). Examination of the value distributions indicate that these positions have very different behaviors regarding amino acid type and variability. For example, position 334 is mostly W (F in Orco); the *A. bakeri* Orco position is 25, in the transmembrane part of the protein. Position 2493 is mostly F, but also significantly Y and L; this is 394 in *A. bakeri* Orco, placed in the transmembrane domain and pointing outside the structure, it could be accessible for phosphorylation, and could indicate a regulatory mechanism. In contrast, other positions have much more variability, like position 1069 or position 1472 (commented above for its association to methyl salicylate and geranyl acetate) corresponding to *A. bakeri* Orco positions 139 and 203, respectively, situated near positions equivalent to experimentally verified OR residues (see **Figure 4C**).

### Relative conservation of predictive residues

To investigate the differential conservation in each cluster of the residues involved in the molecular function with respect to the general conservation of the OR sequence (including regulatory motifs and positions for interaction with other proteins), we annotated each cluster with more than five ORs (101 clusters) with the ratio between the amino acid conservation at the predictive positions (defined as the union of those among the top 10 of the three models; 29 residues, **Table 3**) and the background of amino acid conservation of the entire sequence (see Methods for details; **Supplementary File S4**). We hypothesize that clusters with high values of this ratio, having a ligand-binding pocket more conserved than the background, would detect a narrow collection of odorants, while clusters with lower values of this ratio, having a binding pocket less conserved than background, would detect a wider collection of odorants. The latter could point to the evolutionary adaptation of an OR group with a conserved biological function (e.g. foraging) to diverse odorants (e.g. associated with changes in diet).

While there is a pretty good linear correlation between predictive residue conservation and background sequence conservation (**Figure 5A**), their ratios range from 0.848 for C0 (one of the two large clusters that includes sequences from all 21 species) to 1.149 for C61 (containing 9 ORs in 3 species of Neoptera) with a median value of 0.979 (**Supplementary File S4**). The ant specific C2, representing largely the 9-exon family, has a value of 0.990.

**Figure 5.**
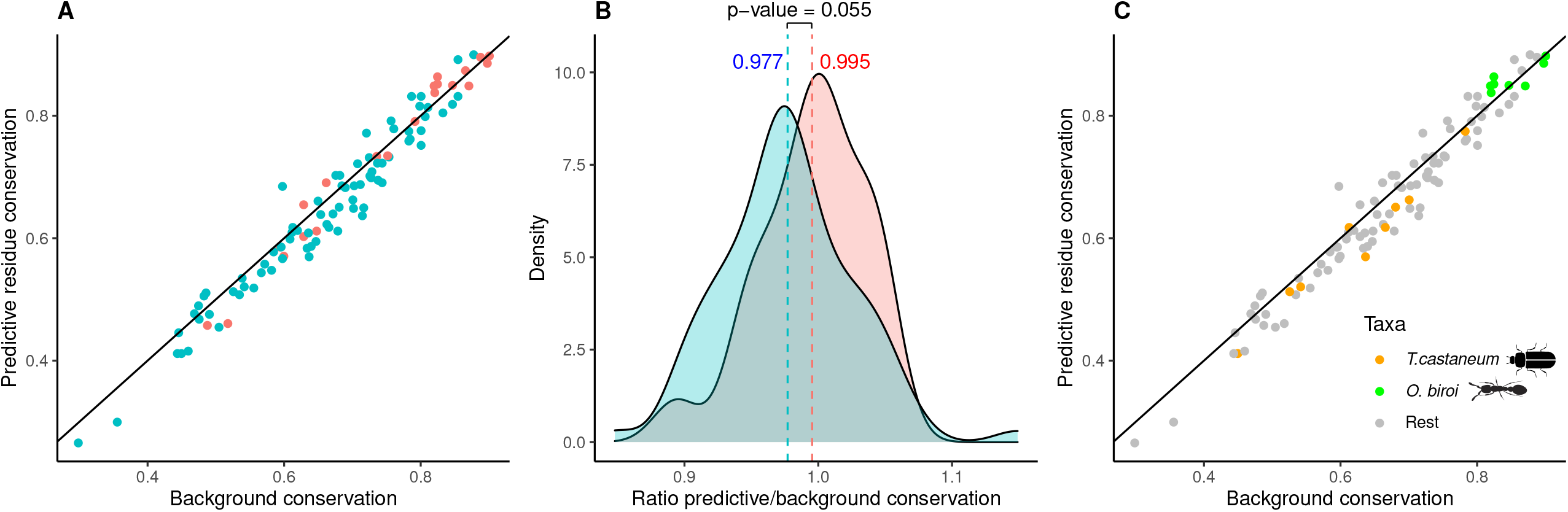
Relative conservation of predictive residues is significantly higher in clusters expanded in social insects. **(A)** Average values of conservation (predictive residues versus the background of the entire sequence) are shown for each cluster. The diagonal indicates clusters where the predictive residues are as conserved as the background. **(B)** Distributions of the values for ratio of predictive to background residue conservation. The 101 clusters with five or more ORs were considered for this analysis (**Supplementary File S4**). Clusters highly expanded in social insects (ratio ORs to species above 3.5 and more than 94% ORs from social insects; 21 clusters; red) have significantly higher relative conservation of predictive residues than other clusters (blue; average values 0.995 and 0.977, respectively; p-value = 0.055, Wilcoxon test). Clusters with many social insects (50 clusters with more than 94% ORs from social insects) do not have a significantly higher relative conservation of predictive residues (p-value = 0.313). Clusters with many species-specific expansions (48 clusters with ratio ORs to species above 3.5) also do not have a significantly higher relative conservation of predictive residues (p-value = 0.529). The thresholds used to segregate the clusters were based on the median values of the respective variable: (i) percentage of ORs from social insects and (ii) ratio ORs to species, respectively. **(C)** Conservation values for clusters from the two species with most species-specific expanded clusters: *T. castaneum* and *O. biroi* (orange and green, respectively; p-value = 0.004).

The numerous OR family expansions within the social insects have been proposed to reflect selective pressure to increase the ability of these species for chemical communication [6,7]. We wondered whether these OR radiations go along with a narrowing or widening of odor tuning, e.g. whether new ORs arising via duplication in a cluster are likely to bind to very similar or dissimilar ligands. Evidence for this would be a larger or lower relative conservation of residues predictive for binding, respectively.

To test this hypothesis, we segregated the 101 clusters in two ways: (i) we defined clusters rich in ORs from social insects as those that had more ORs from social insects than the median content of all clusters (94%; 50 clusters), and (ii) we defined highly expanded clusters as those that had a ratio of ORs to species represented in the cluster higher than the median (3.5 ORs per species; 48 clusters). Segregating the clusters by either condition (i) or (ii) did not result in significantly different distributions of relative conservation of predictive residues (p-values of 0.313 and 0.529, respectively, Wilcoxon test). Strikingly, we observed a more significant result when both conditions were applied together (21 clusters; p-value = 0.055; **Figure 5B**), which suggests that clusters with many expansions in the social insects have indeed a trend towards higher relative conservation of these residues. Social evolution in the insects is therefore characterized by duplications of genes leading to large OR subfamilies specialized in the detection of very similar odorants.

Focusing on the two insects from our set of 21 species with the highest number of species-specific expanded clusters, the beetle *T. castaneum* (9 clusters in **Figure 5C**; orange), a non-social insect, and the ant *O. biroi* (8 clusters in **Figure 5C**; green), a social insect, we note that OR clusters from the beetle exhibit a lower level of residue conservation than the ant (0.956 and 1.014, respectively; p-value = 0.004). These numbers suggest that the evolutionary and functional processes at play shaping the evolution of the OR family must be different between these species. Higher OR expansion more involved in adaptation to novel odorants in the beetle *T. castaneum* than in the ant *O. biroi*, and more regulatory tuning of ORs detecting narrowly defined odorants in the ant than in the beetle, are evolutionary hypotheses consistent with our observations.

## Conclusions

In this work we have presented a novel approach to expand the functional characterization of the large OR protein family using machine learning. Our approach starts with the collection and curation of selected datasets of insect ORs. The alignment of the 3,902 protein sequences provides a framework to compare functional information from these sequences. Positions in this alignment were mapped to a template structure, which is available for one ancestral protein of the family [30]. We used then machine learning to study three separate functionally characterized datasets, each of them providing sets of residues predicted to determine ligand recognition. Independently, we used a sequence-based clustering algorithm to separate the family in clusters, expected to be responsible for related functions in the same or in different organisms. Finally, we annotated these clusters for their taxonomic distribution, identifying clusters with particular expansion patterns, and with different relative conservation of residues predicted to be associated to ligand recognition. Our results suggest that the large expansions of the OR family in social insects are associated to subfamilies detecting very similar ligands (**Figure 5**). Expansions leading to subfamilies with wider detection ranges might be more exclusive to non-social species like *T. castaneum*.

Our work facilitates the analysis of ORs from 21 insect species with respect to information that we obtained for the entire family. These data are available through a dedicated web service called iOrME. Potentially, other individual ORs from species not included in our set of 21 species can be added by incorporating them to the multiple sequence alignment of 3,902 sequences. In this sense, all mapped information can flow from and to new OR sequences of interest.

We are aware that the results presented in our work must be necessarily affected by the choice of species, which reflects itself a bias in this field of research, but we have tried to remove such biases by defining variables that can be applied to different taxonomic levels and that are normalized by values like number of species or conservation of entire sequences. As part of an effort to remove these biases, we plan to add novel OR datasets as required, and in principle it should be easy to include also new experimental data and information from new protein structures as they become available. Our dedicated web site constitutes a resource that will accommodate newer versions of the OR dataset, clusters, machine learning results and annotations.

The OR family is not the only large protein family with large paralogous expansions (see e.g. ubiquitination-related families in Chlamydiae [52], or the families of F-box proteins in plants [53]). We propose that an approach like the one we have presented here could be similarly applied to other expanded families, irrespective of their function or taxonomic distribution. We expect that from the study of many such families, we will obtain further insights into the rules that drive gene duplication and gain of function.

## Data Availability

The datasets used, produced and analyzed in our study are available in the iOrME repository (http://cbdm-01.zdv.uni-mainz.de/~munoz/iorme/).

## Acknowledgements

We thank Mohamed Kamel (University of Bejaia, Algeria) for assistance in sequence clustering and Hugh M. Robertson (University of Illinois Urbana-Champaign) for help in the collection of OR protein sequence datasets.

## Supplementary Data

**Figure S1**: Number of proteins in each cluster for subfamilies defined by gene models for species *A. cephalotes, A. echinatior, C. floridanus* and *H. saltator*.

**Figure S2**: Average F1, precision and sensitivity values during 10-fold cross-validations repeated 10 times of the machine learning models (see Methods for details).

**File S1**: Multiple sequence alignment of all 3,902 OR sequences considered in the analysis.

**File S2**: Binarized response data of OR proteins from 3 datasets to 8 chemicals. Columns indicate: dataset (#1 = *D. melanogaster*, #2 = *A. gambiae*, #3 = *H. saltator*), id - cluster identifier, ac - UniProt AC, following eight columns - registry numbers of the panel of chemicals (see Methods for IUPAC and common names).

**File S3**: Clusters of OR proteins.

**File S4**: Annotated clusters. Columns indicate: ID - cluster ID, Number OR - number of ORs, Protein length - average length with standard deviation, Number of species, Common taxonomy, Bg conservation - conservation of the entire sequence, Predictive conservation – conservation of predictive residues, Ratio predictive/bg - ratio between the conservation of predictive residues and the entire sequence, Social ratio – ratio of ORs from social insects, ORs/species – ratio between the number of ORs and the number of species in the cluster. See Methods for details.

**File S5**: Classification performance of the machine learning models (a model for each chemical- dataset pair). Columns indicate: dataset (#1 = *D. melanogaster*, #2 = *A. gambiae*, #3 = *H. saltator*), chem_id - chemical ID, mtry - number of variables in each tree of the forest (random forest parameter), auc - area under roc curve, auc_ci - auc confidence interval, sen - sensitivity, sen_ci - sensitivity confidence interval, pre - precision, pre_ci - precision confidence interval, f1 - F1 performance.

**File S6**: Importance of machine learning features (alignment position and amino acid) by model (a model for each chemical-dataset pair). Columns indicate: importance, auc - area under ROC curve of the related model, dataset (#1 = *D. melanogaster*, #2 = *A. gambiae*, #3 = *H. saltator*), chem_id - chemical ID, position - position in the multiple sequence alignment, aa - amino acid.

**File S7**: Amino acid usage for each of the predictive positions in clusters.

**File S8**: Mapping of the positions from the OR alignment to the Orco proteins from *D. melanogaster* and *A. bakeri*.

