## Supplementary figures and images for "Identification of functional residues using machine learning provides insights into the evolution of odorant receptor gene families in solitary and social insects"

### Supplementary Figure S1

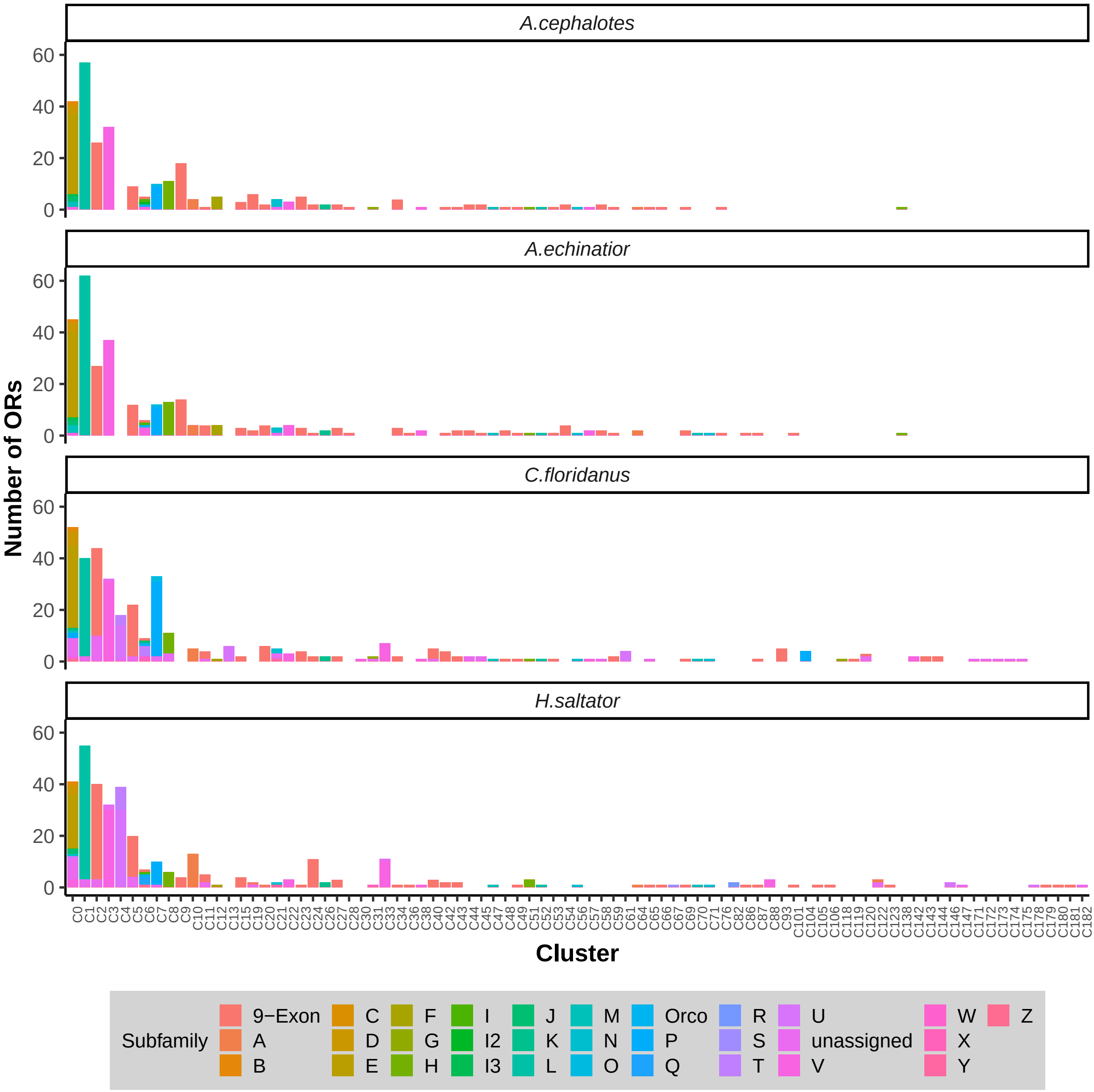

### Supplementary Figure S2

#### Cross-Validation Performance

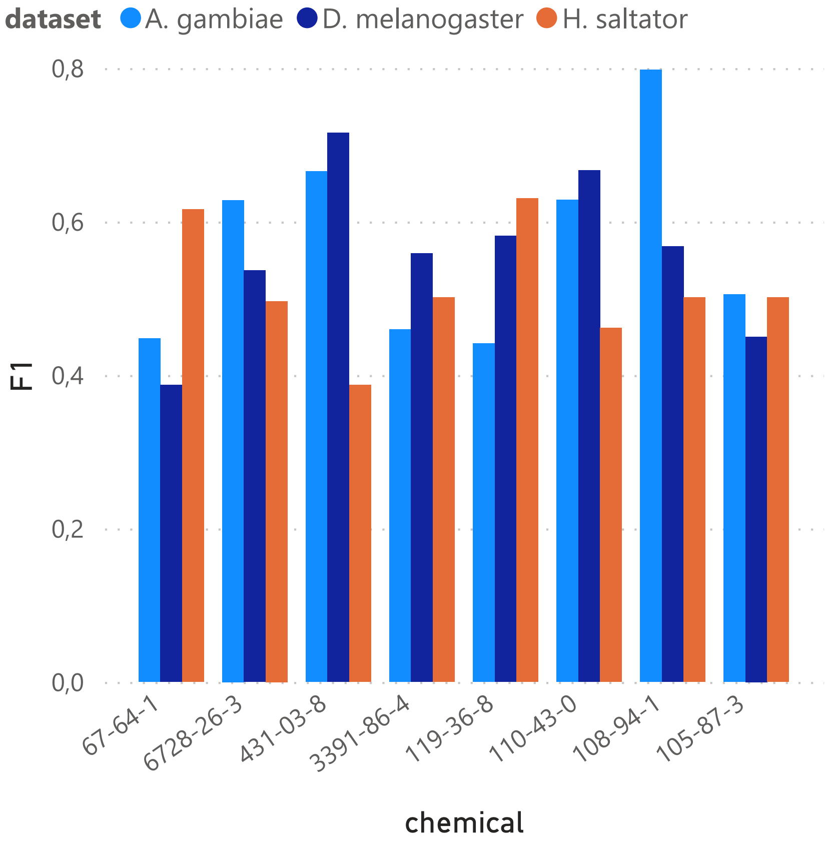


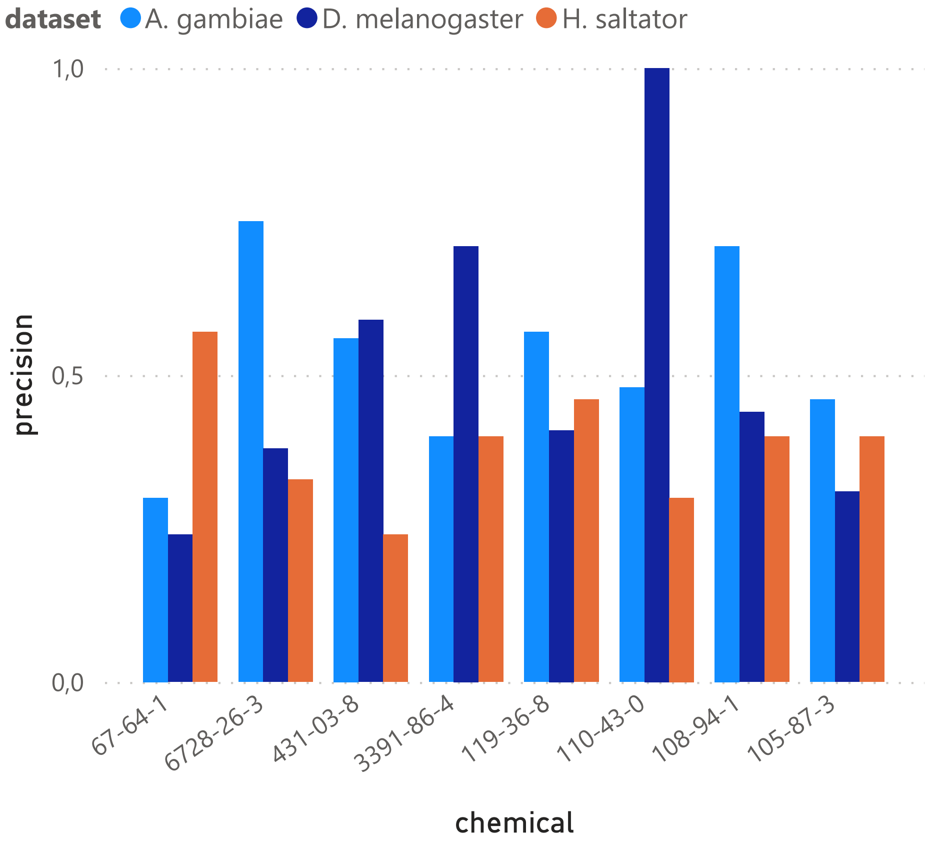


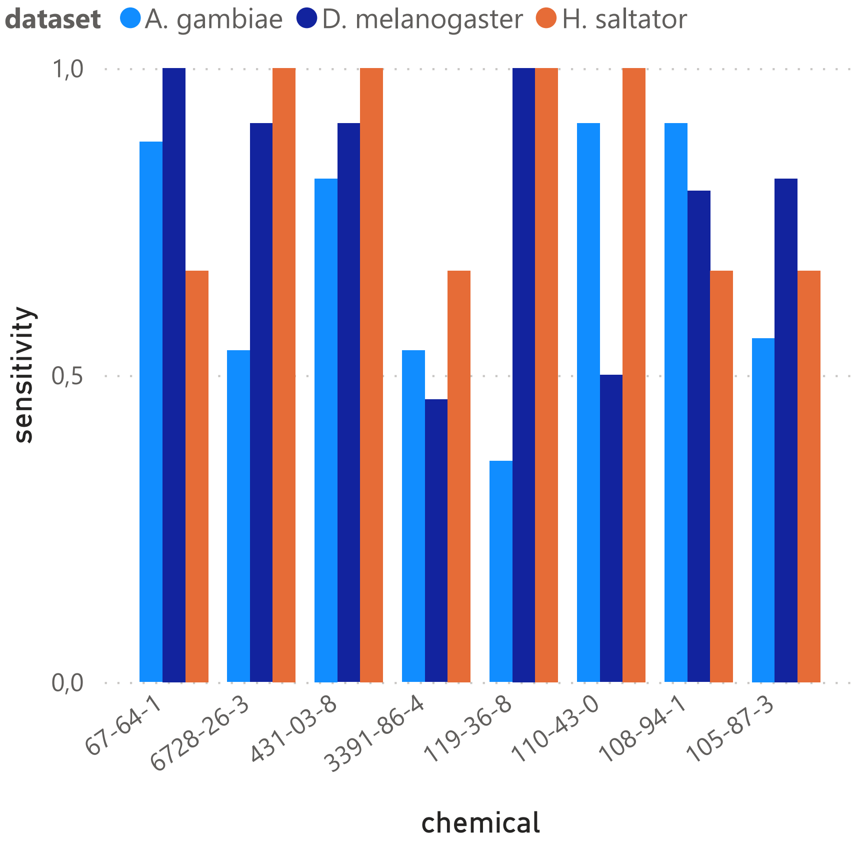
