## Supplementary File S7 for "Identification of functional residues using machine learning provides insights into the evolution of odorant receptor gene families in solitary and social insects"

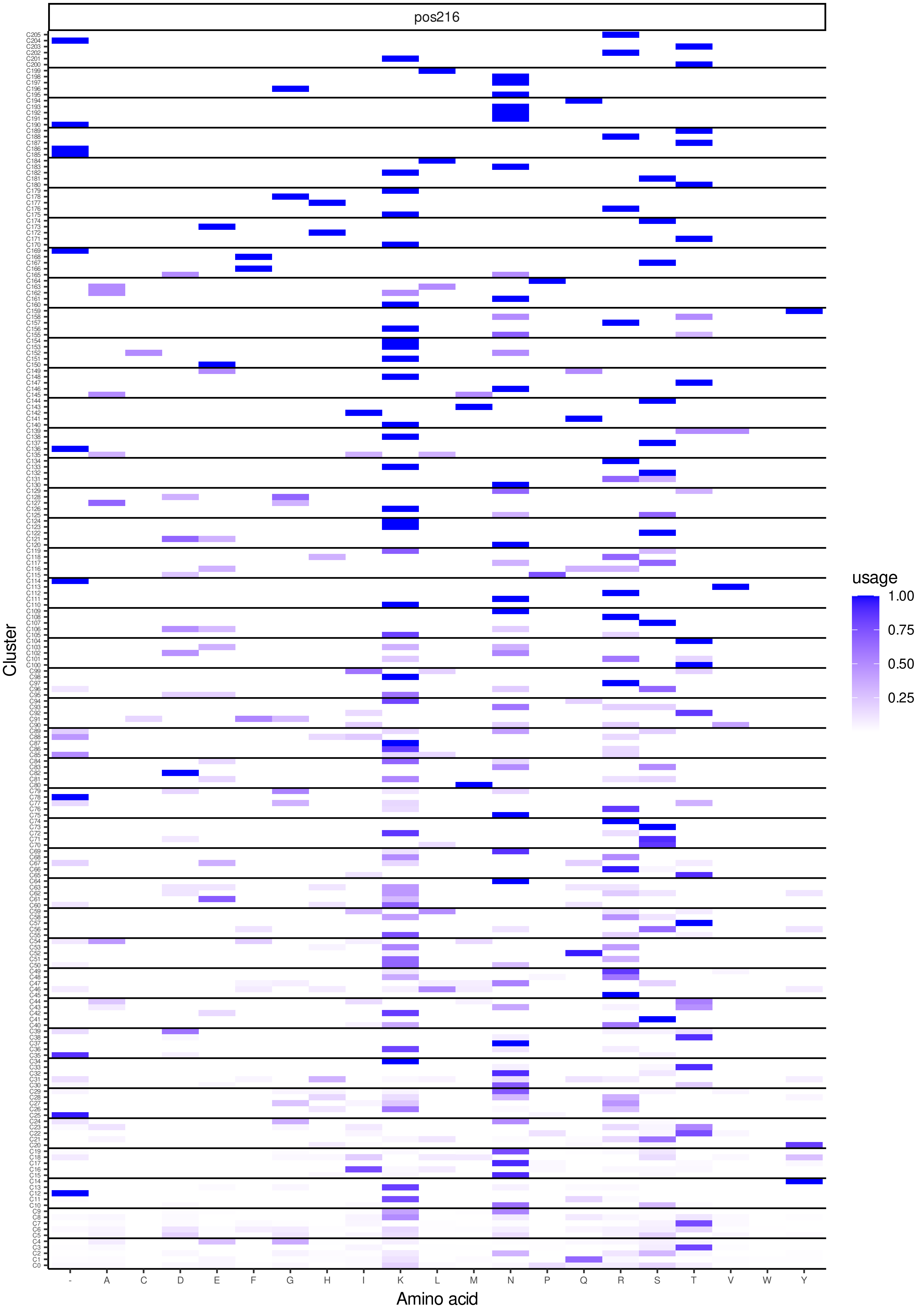
pos216
Cluster
Amino acid
- A C D E F G H I K L M N P Q R S T V W Y
usage
1.00
0.75
0.50
0.25

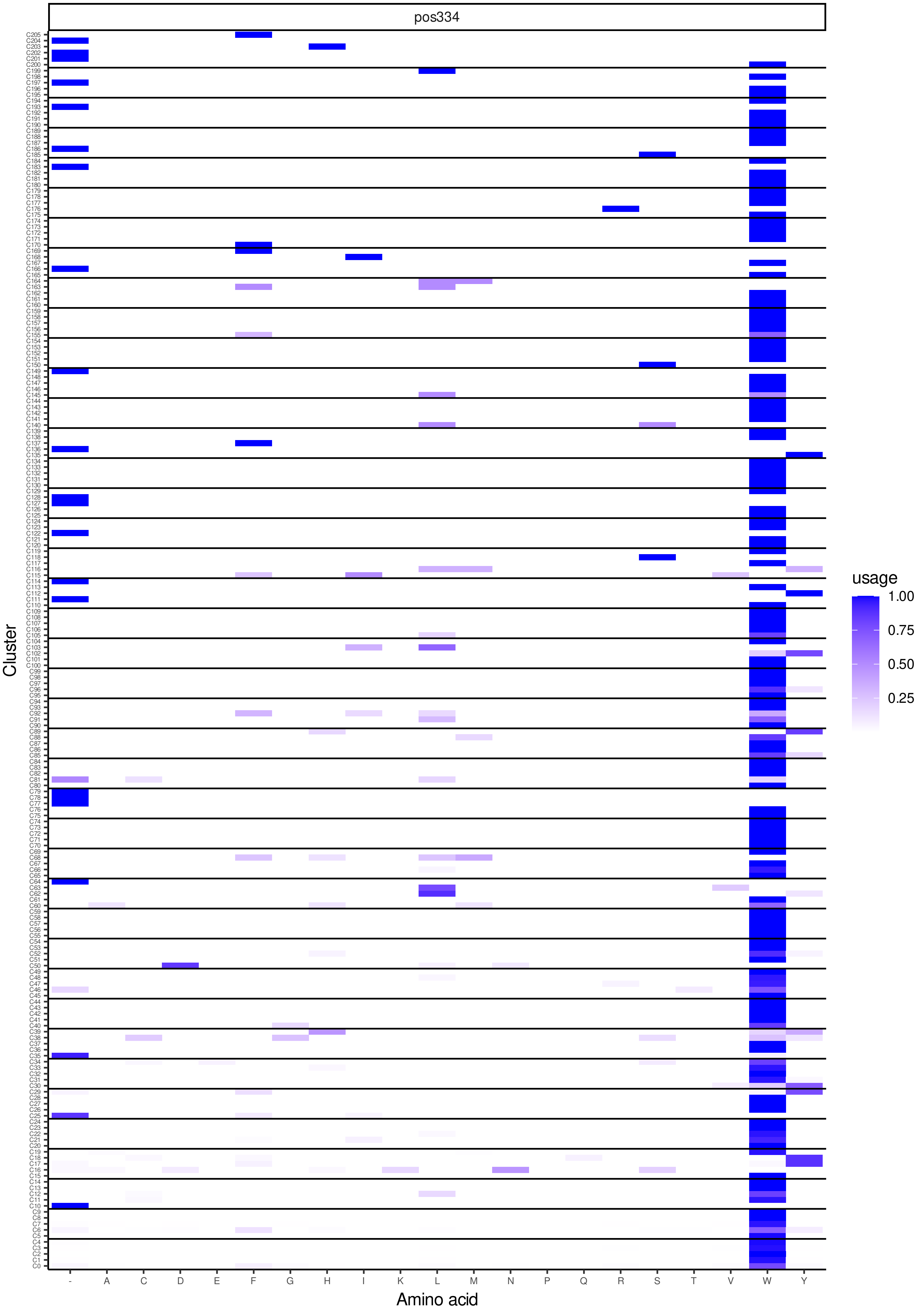
pos334
Cluster
Amino acid
usage
1.00
0.75
0.50
0.25
- A C D E F G H I K L M N P Q R S T V W Y

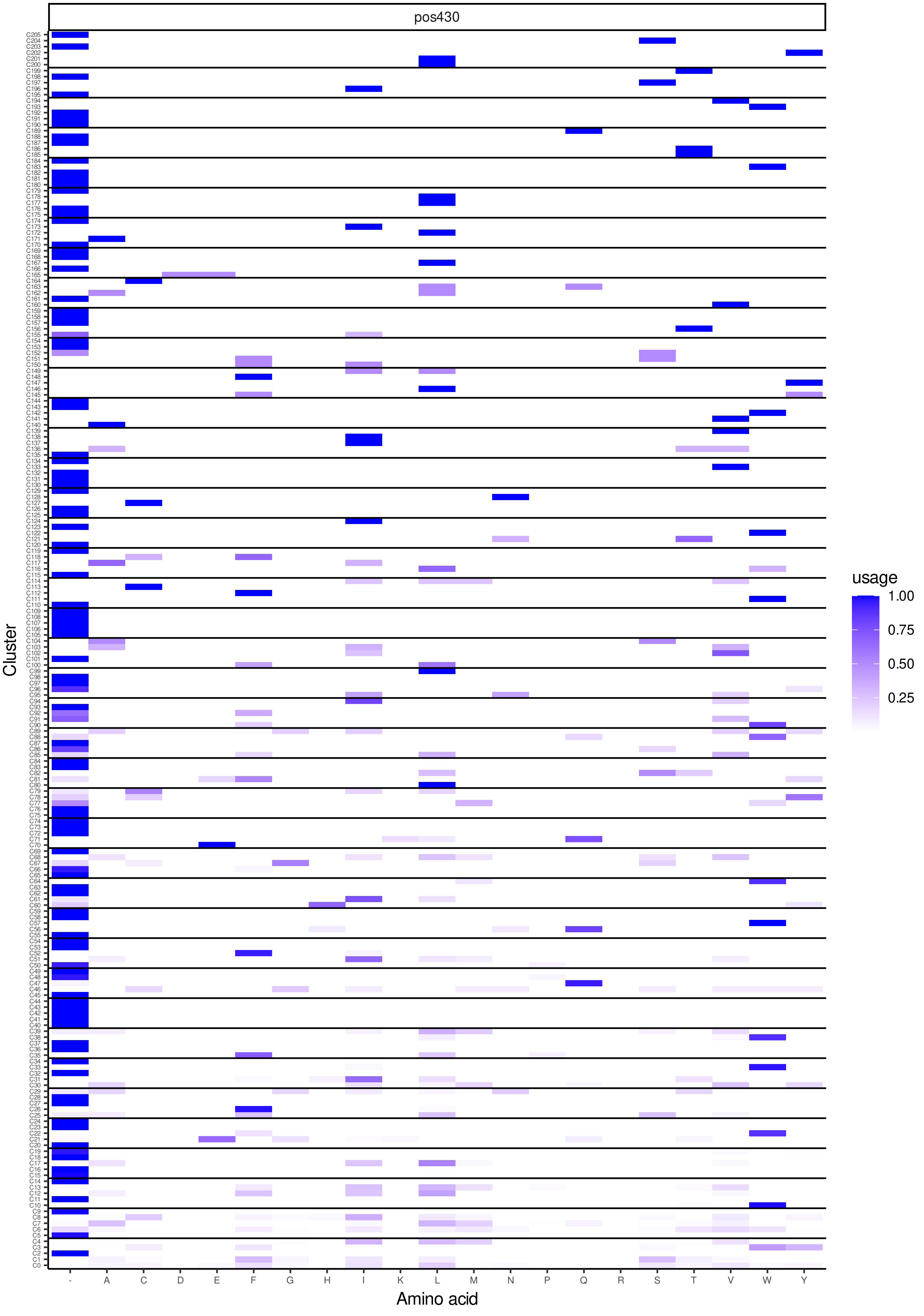
pos430
Cluster
Amino acid
usage
1.00
0.75
0.50
0.25
- A C D E F G H I K L M N P Q R S T V W Y

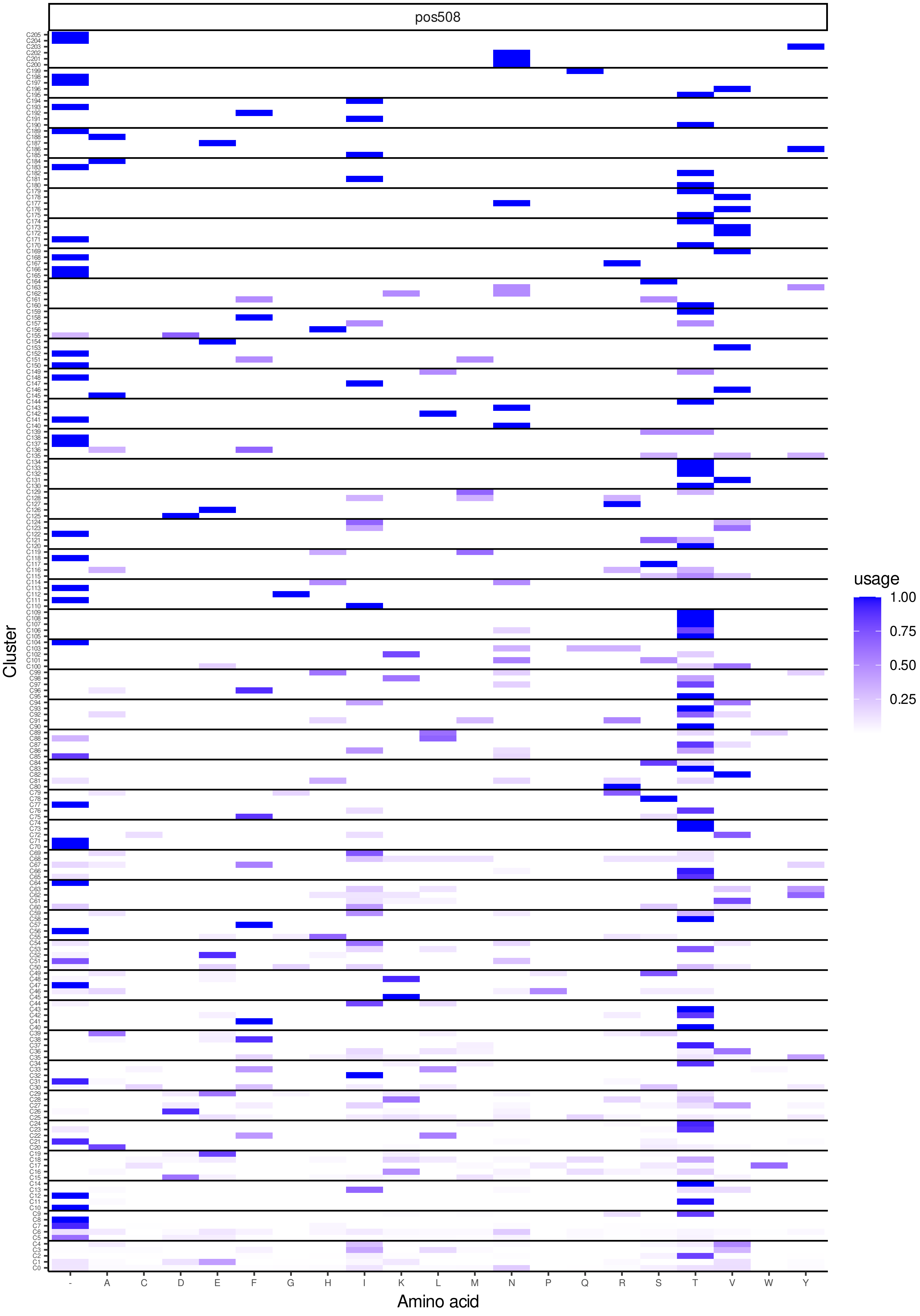
pos508
Cluster
Amino acid
- A C D E F G H I K L M N P Q R S T V W Y
C205 … C0
usage
1.00
0.75
0.50
0.25

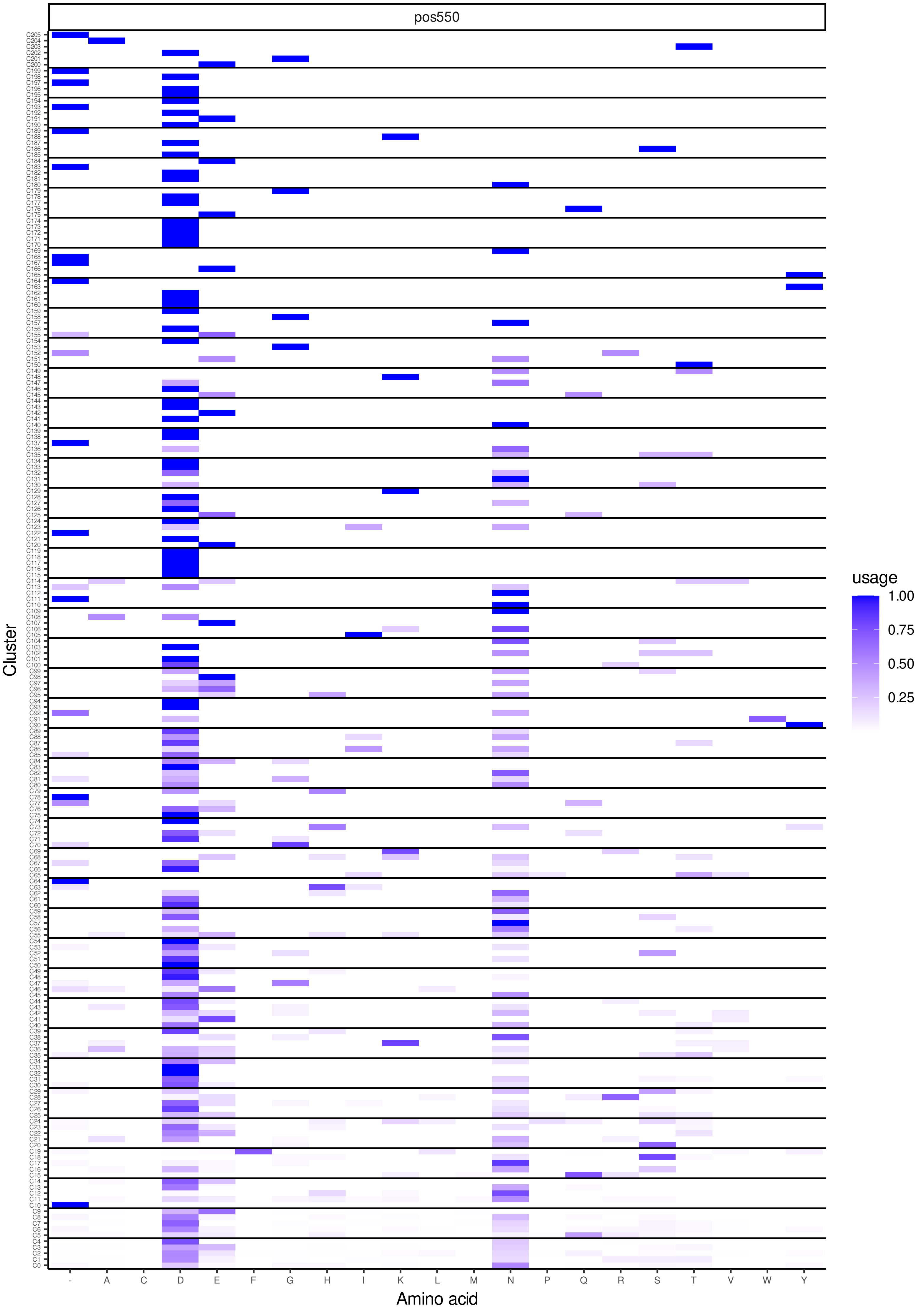
pos550
usage
1.00
0.75
0.50
0.25
Cluster
Amino acid
- A C D E F G H I K L M N P Q R S T V W Y

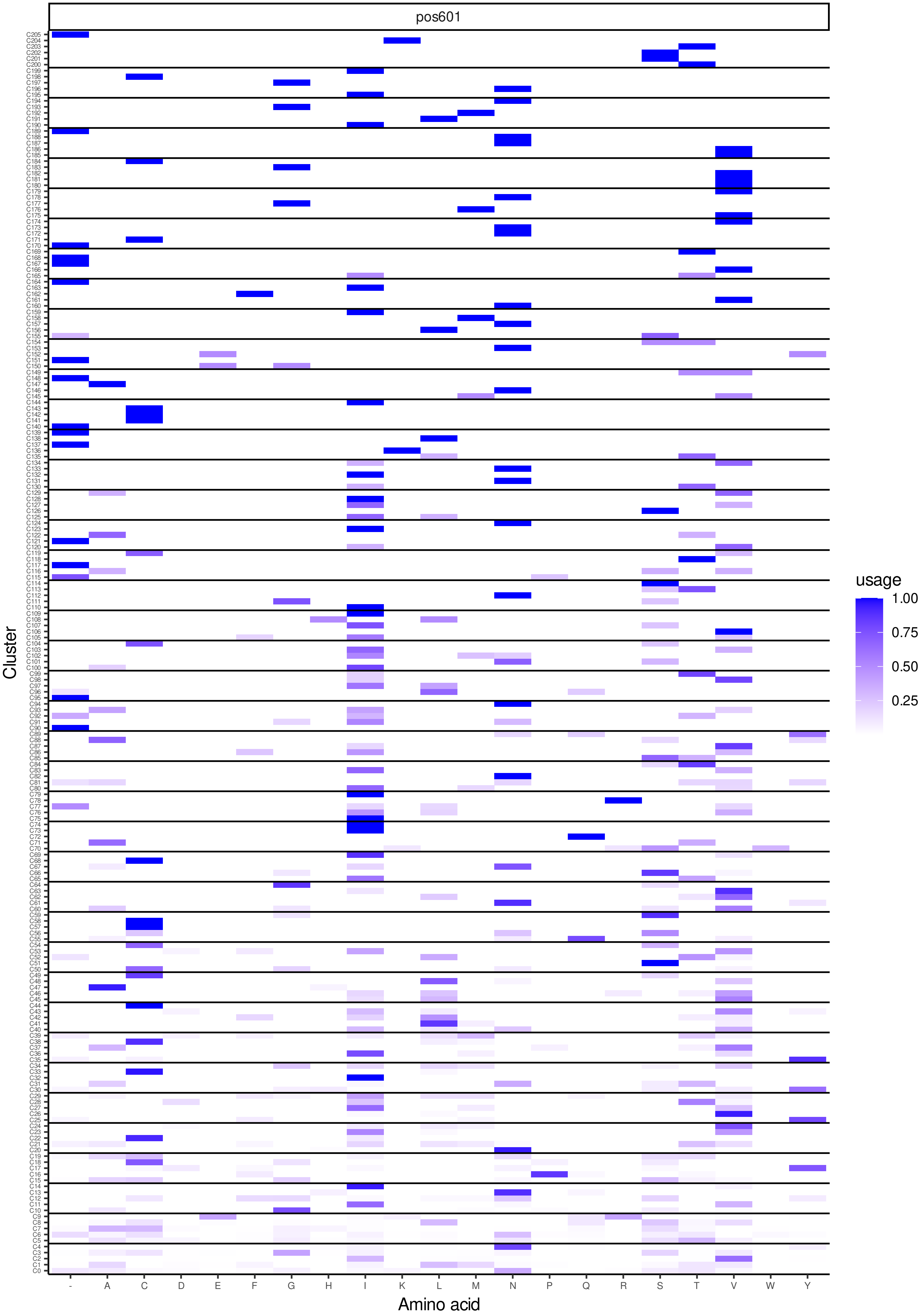
pos601
usage
1.00
0.75
0.50
0.25
Cluster
Amino acid
- A C D E F G H I K L M N P Q R S T V W Y

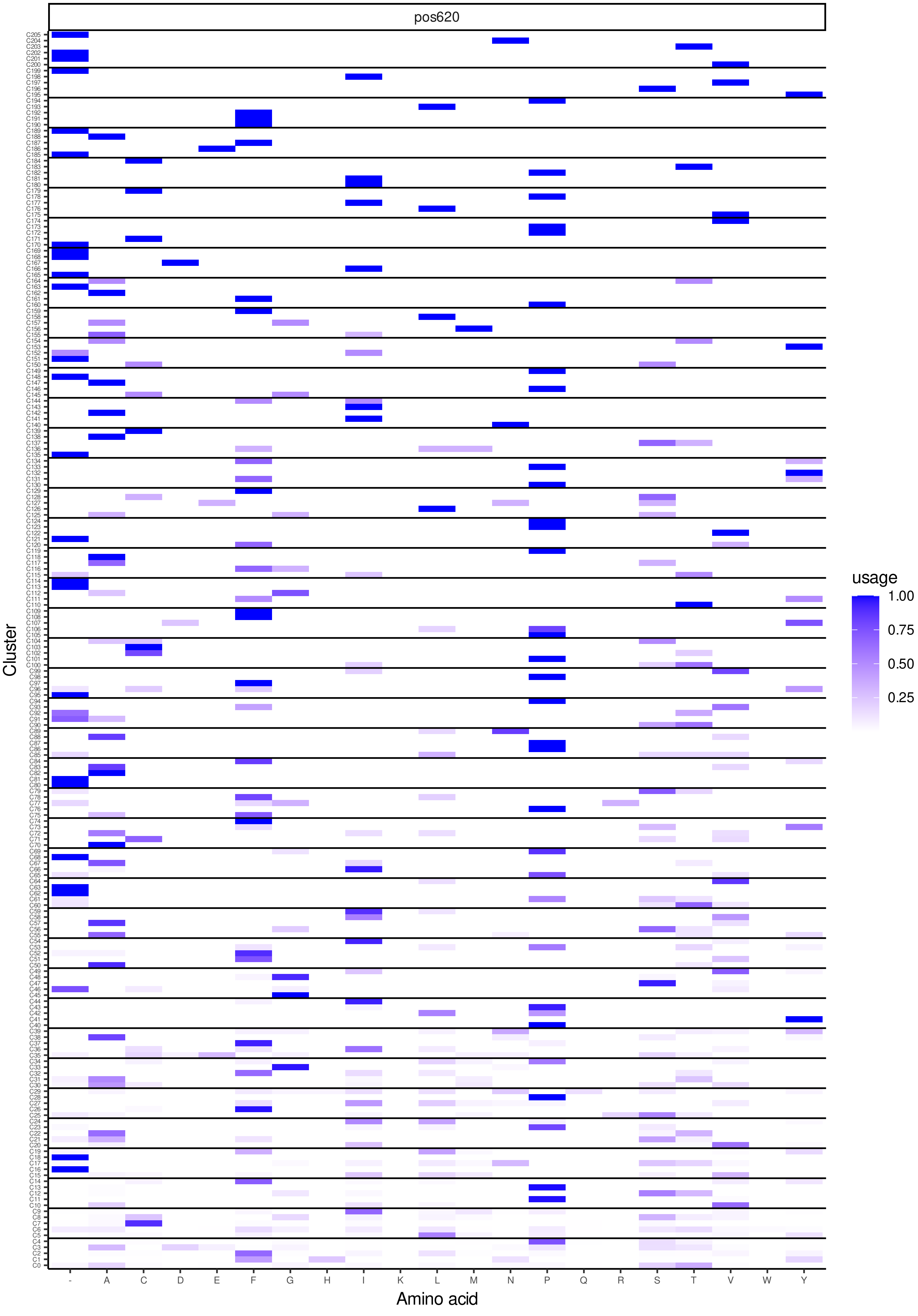
pos620
Cluster
Amino acid
usage
1.00
0.75
0.50
0.25
- A C D E F G H I K L M N P Q R S T V W Y

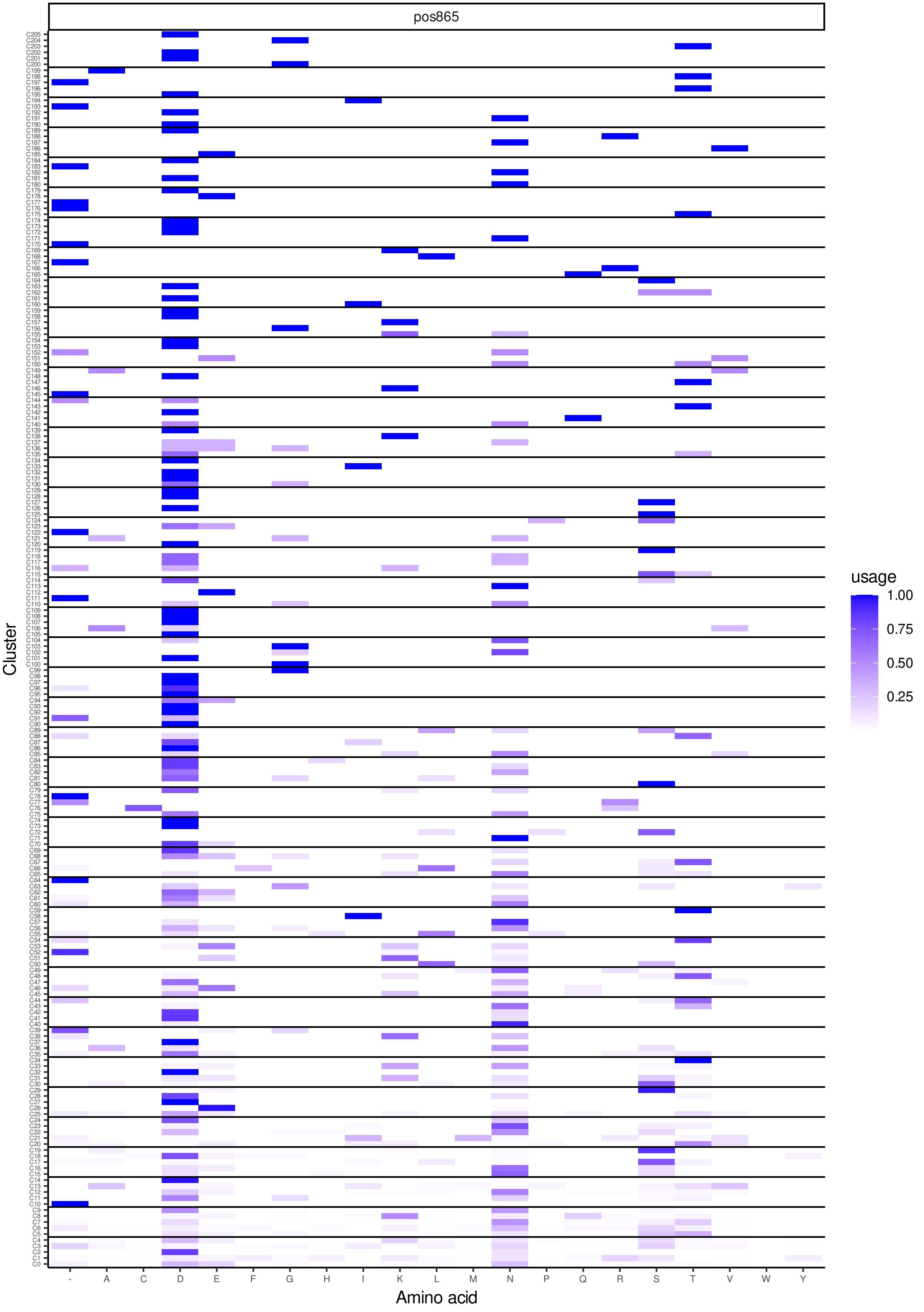
pos865
Cluster
- A C D E F G H I K L M N P Q R S T V W Y
Amino acid
usage
1.00
0.75
0.50
0.25

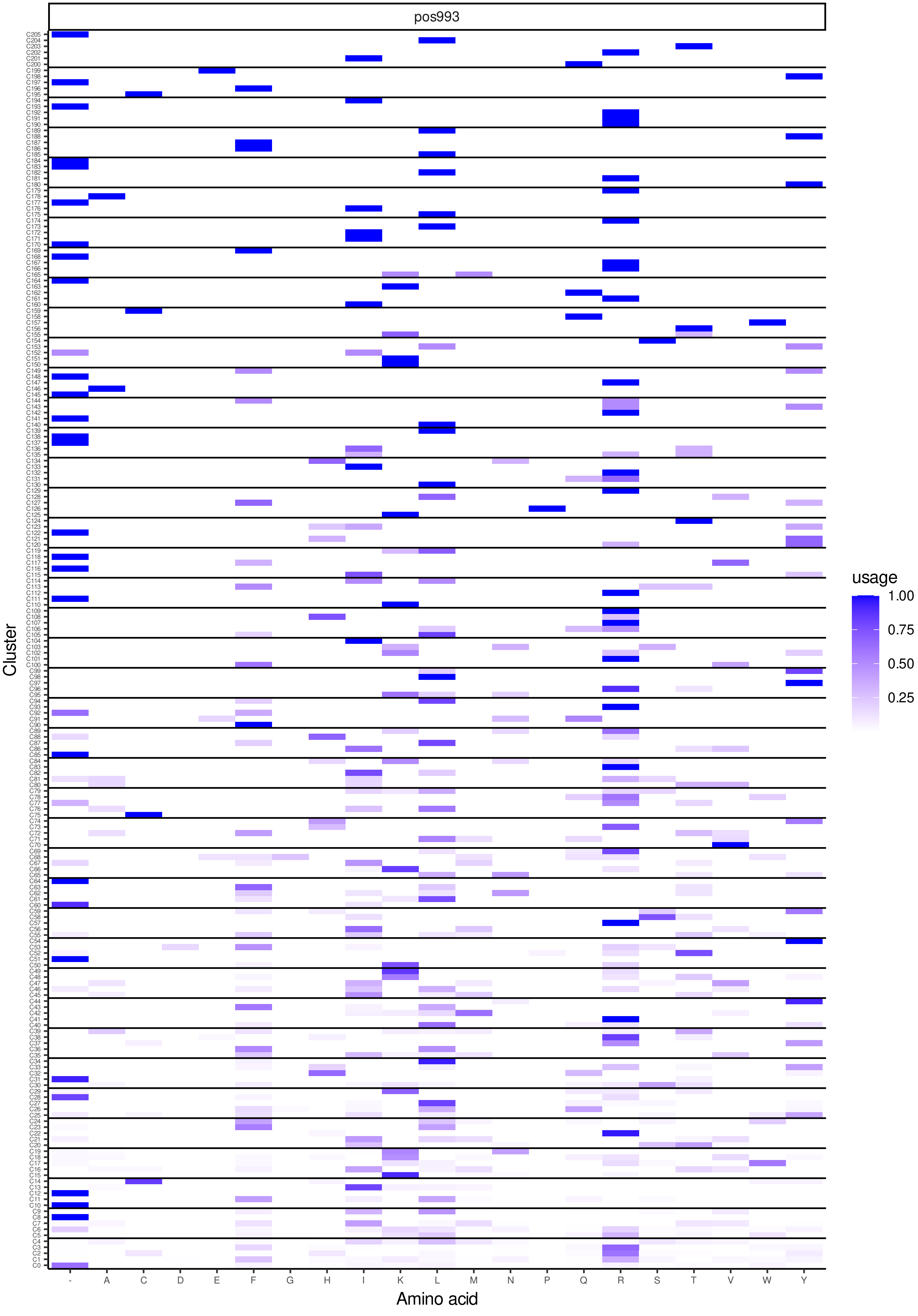
pos993
Cluster
Amino acid
- A C D E F G H I K L M N P Q R S T V W Y
usage
1.00
0.75
0.50
0.25

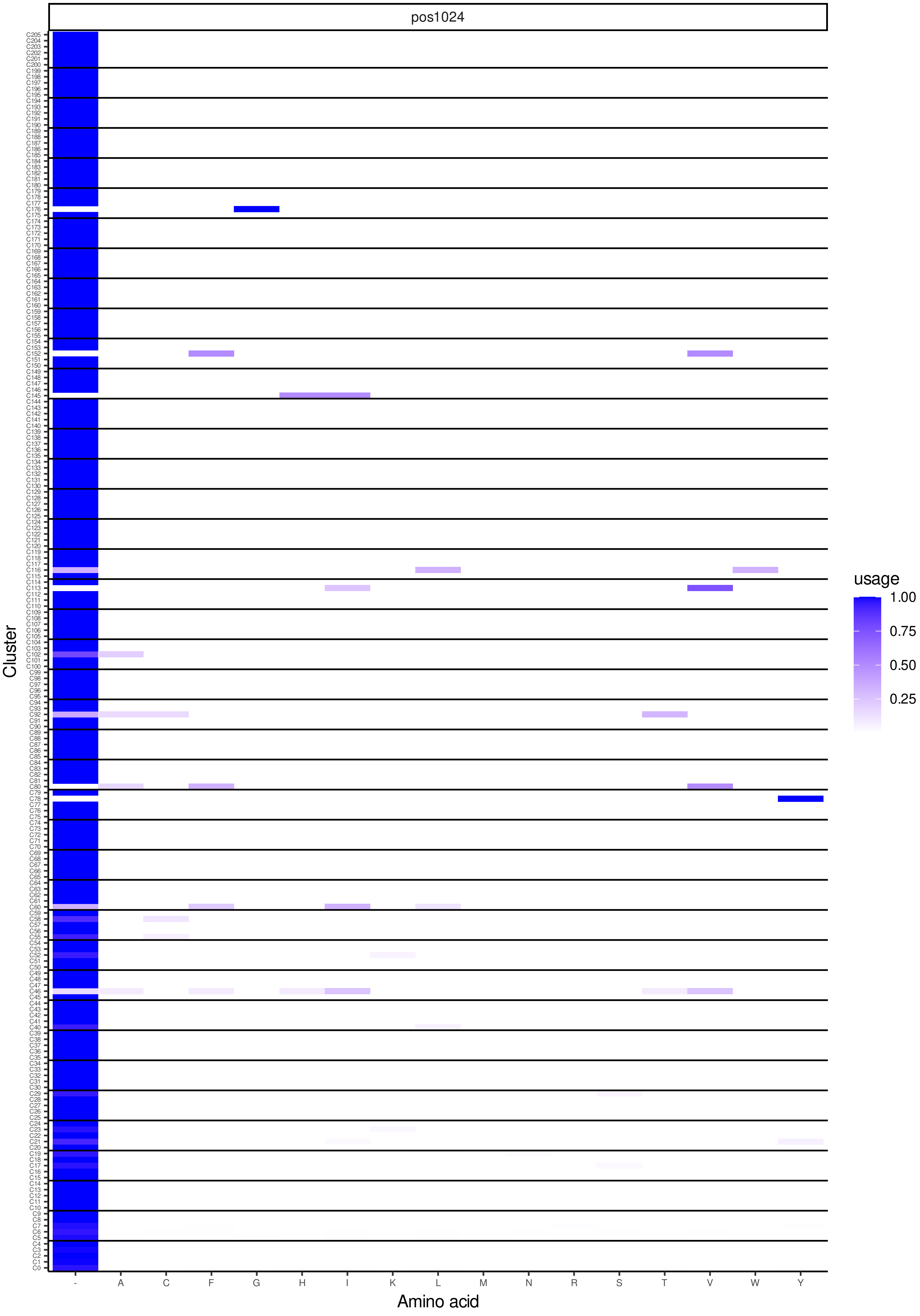
pos1024
Cluster
Amino acid
usage
1.00
0.75
0.50
0.25
- A C F G H I K L M N R S T V W Y

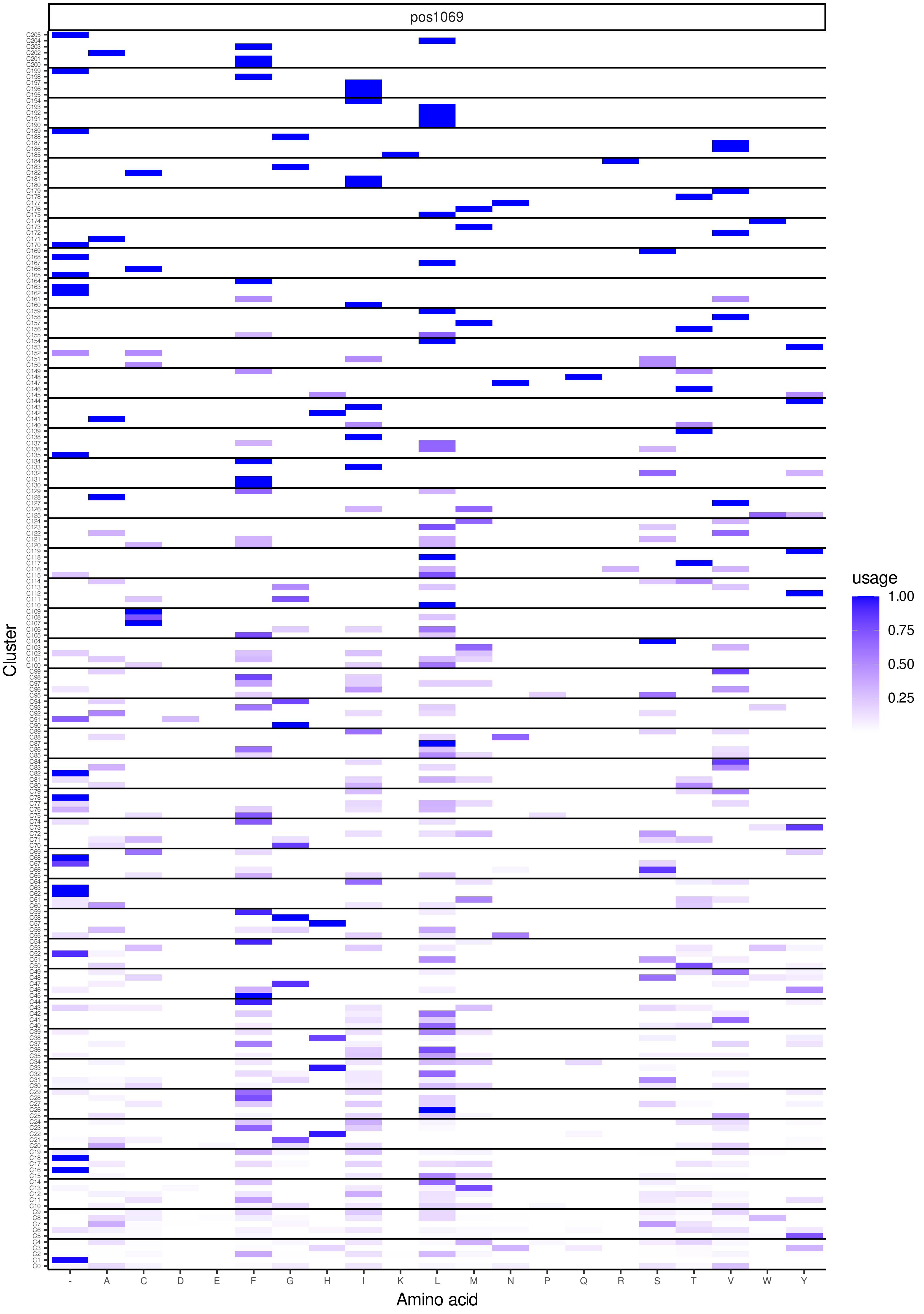
pos1069
Cluster
Amino acid
- A C D E F G H I K L M N P Q R S T V W Y
usage
1.00
0.75
0.50
0.25

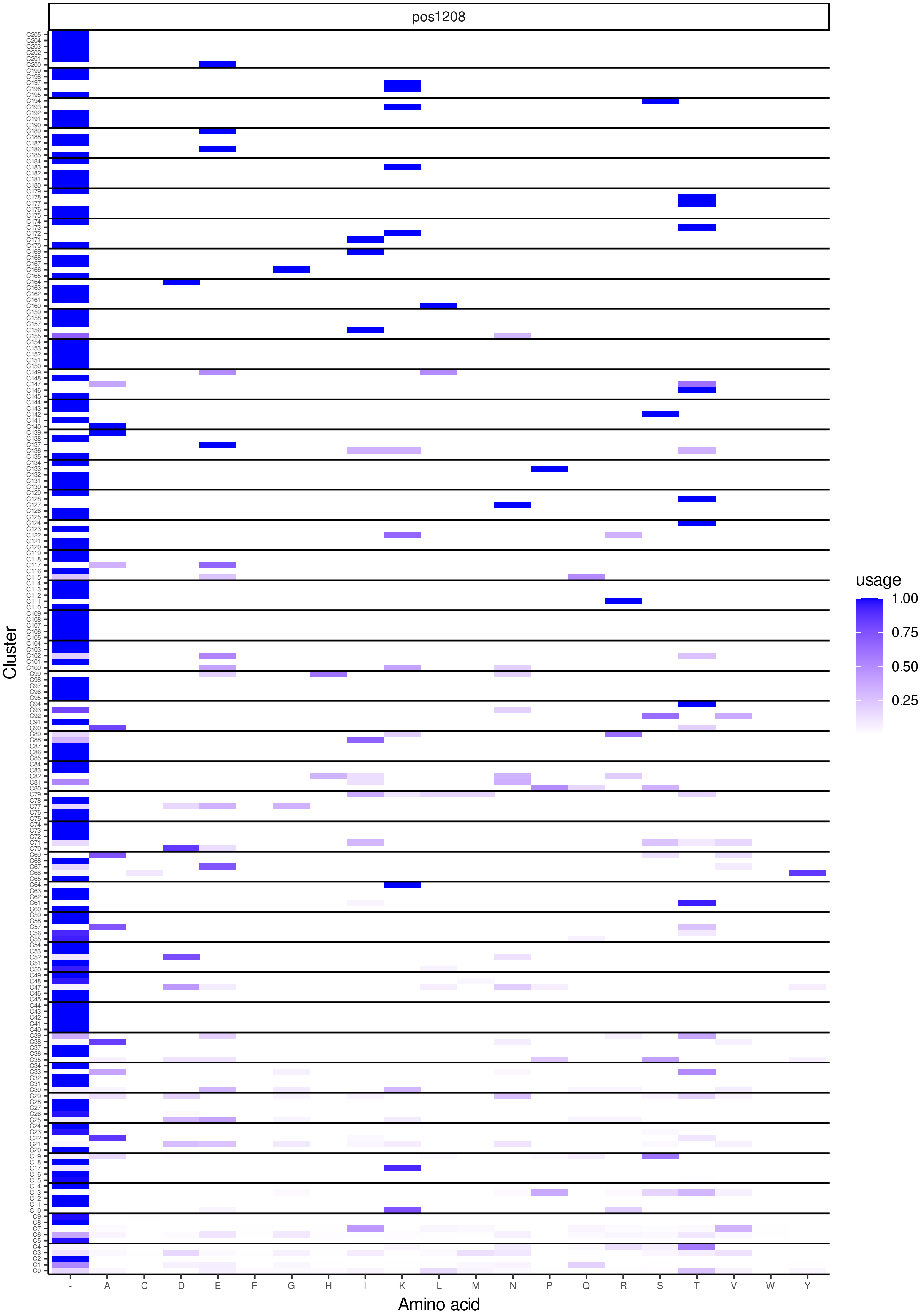
pos1208
Cluster
Amino acid
usage
1.00
0.75
0.50
0.25
- A C D E F G H I K L M N P Q R S T V W Y

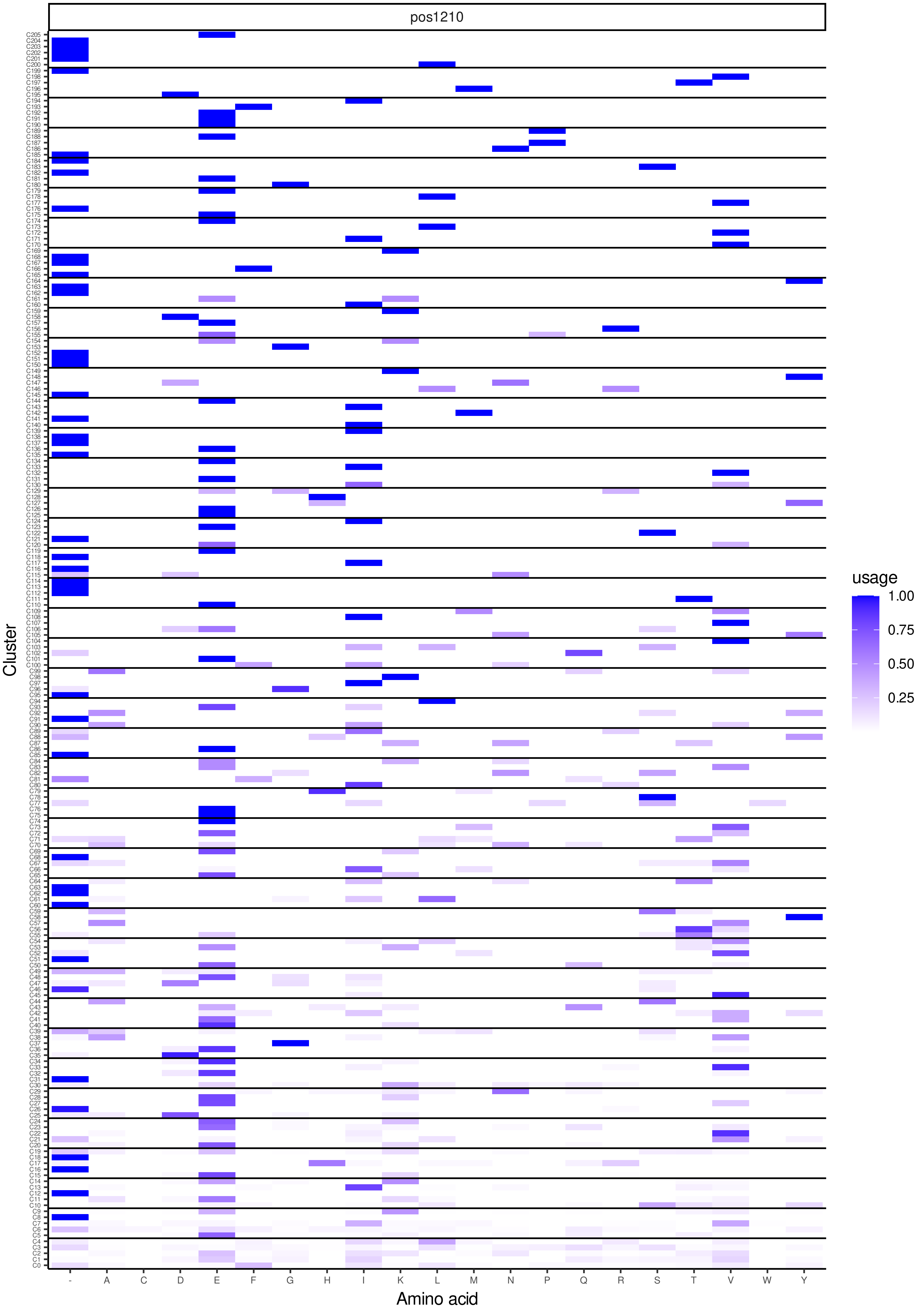
pos1210
Cluster
Amino acid
- A C D E F G H I K L M N P Q R S T V W Y
usage
1.00
0.75
0.50
0.25

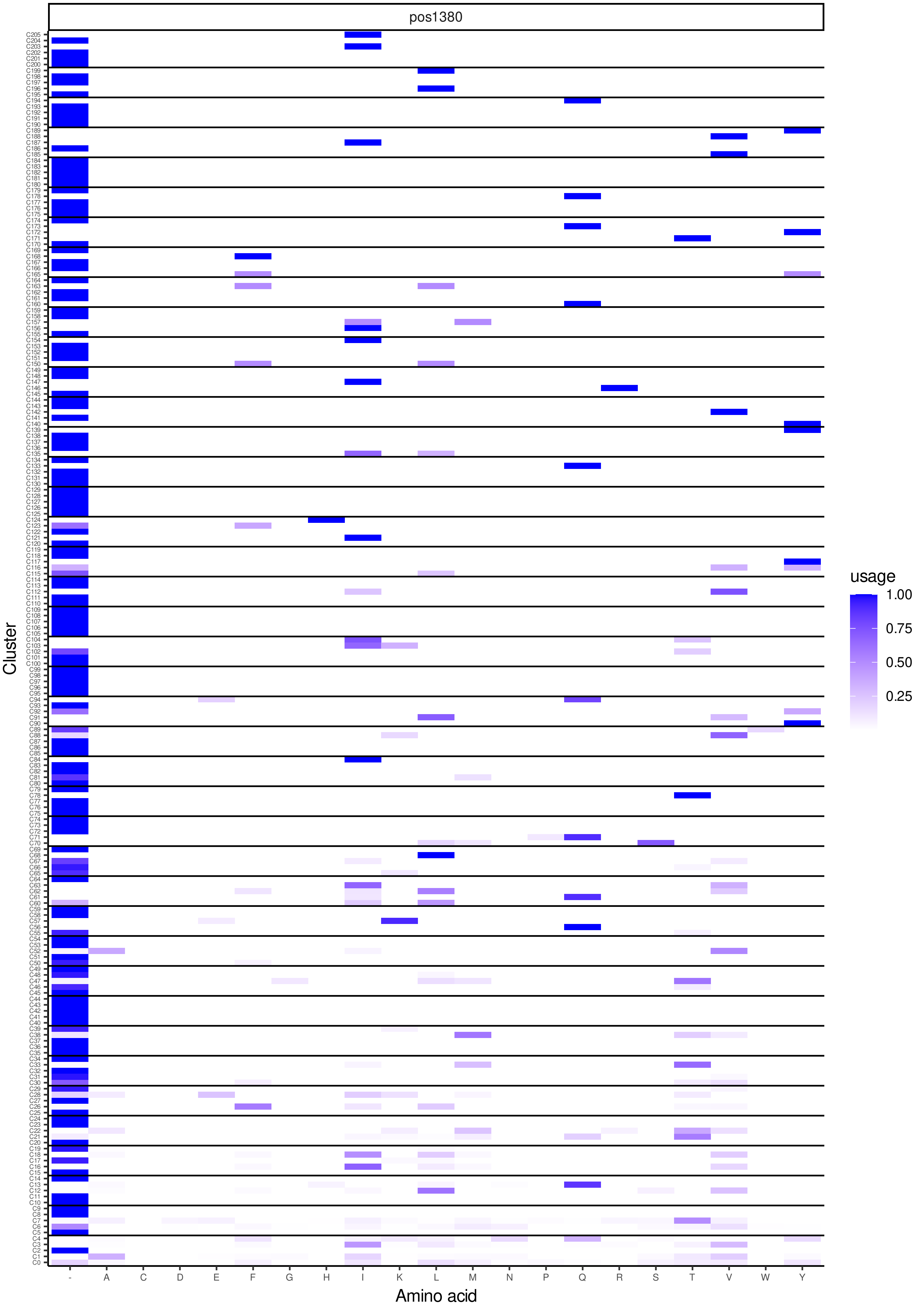
pos1380
Cluster
Amino acid
usage
1.00
0.75
0.50
0.25
- A C D E F G H I K L M N P Q R S T V W Y

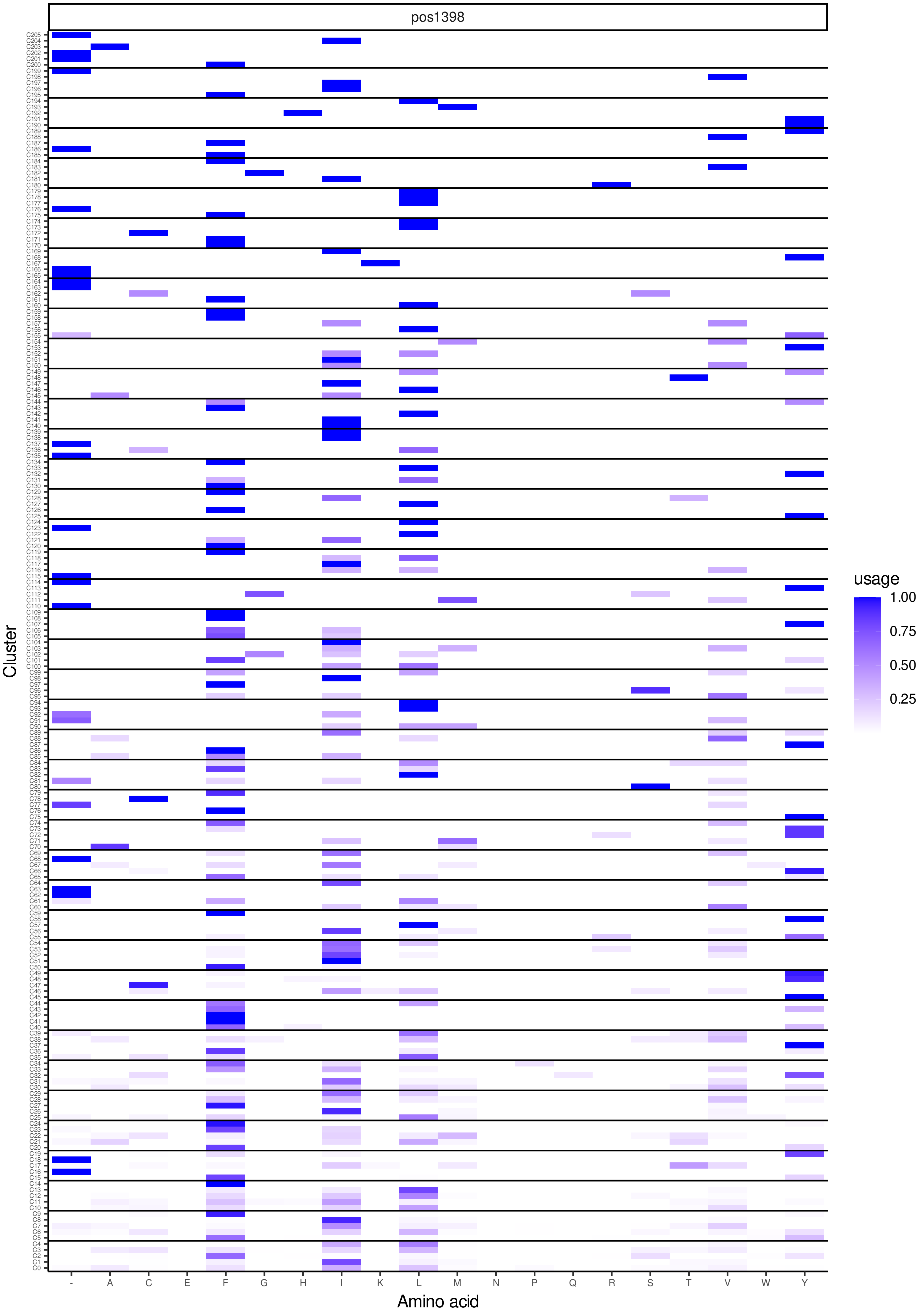
pos1398
Cluster
Amino acid
usage
1.00
0.75
0.50
0.25
- A C E F G H I K L M N P Q R S T V W Y

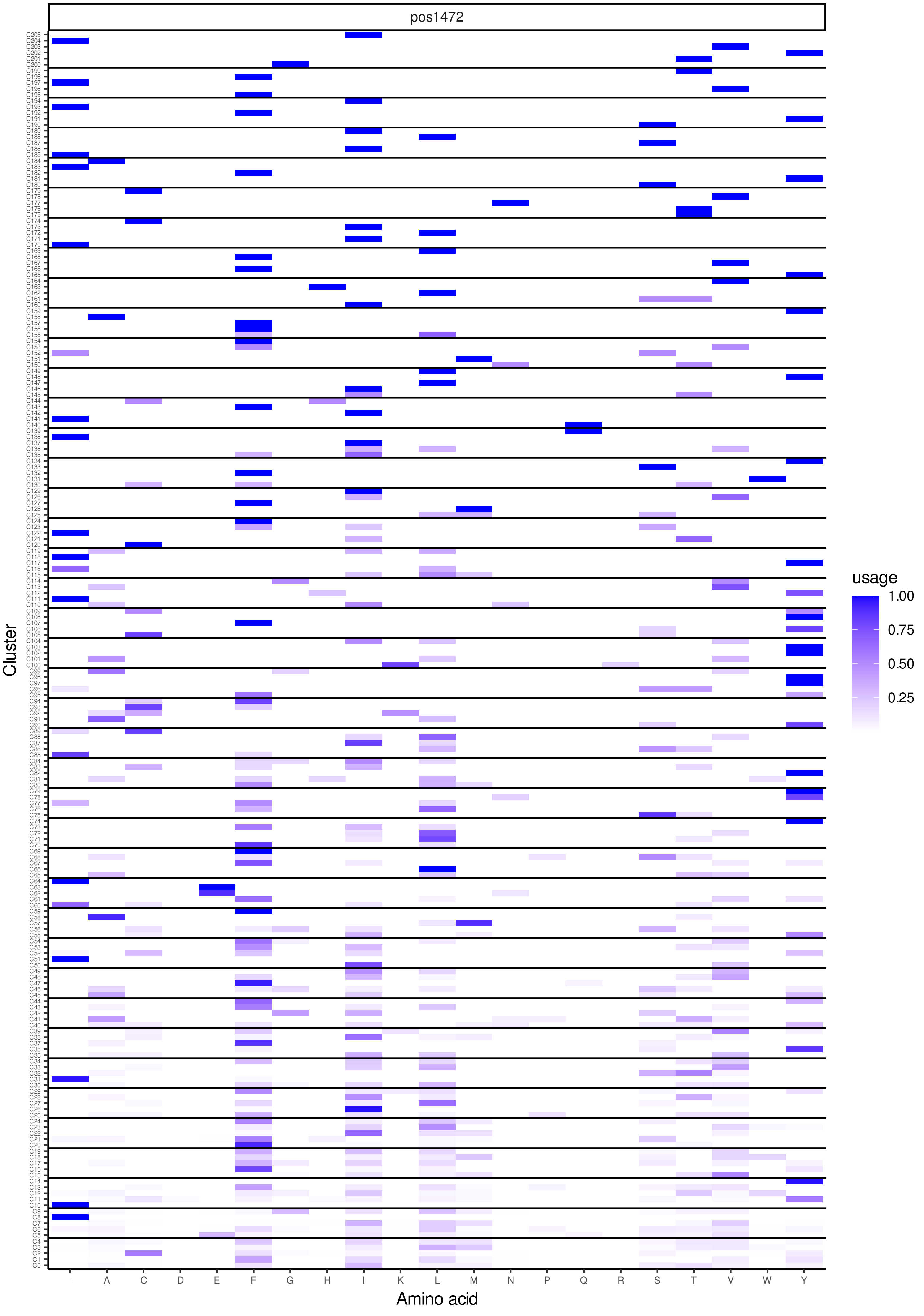
pos1472
Cluster
Amino acid
- A C D E F G H I K L M N P Q R S T V W Y
usage
1.00
0.75
0.50
0.25

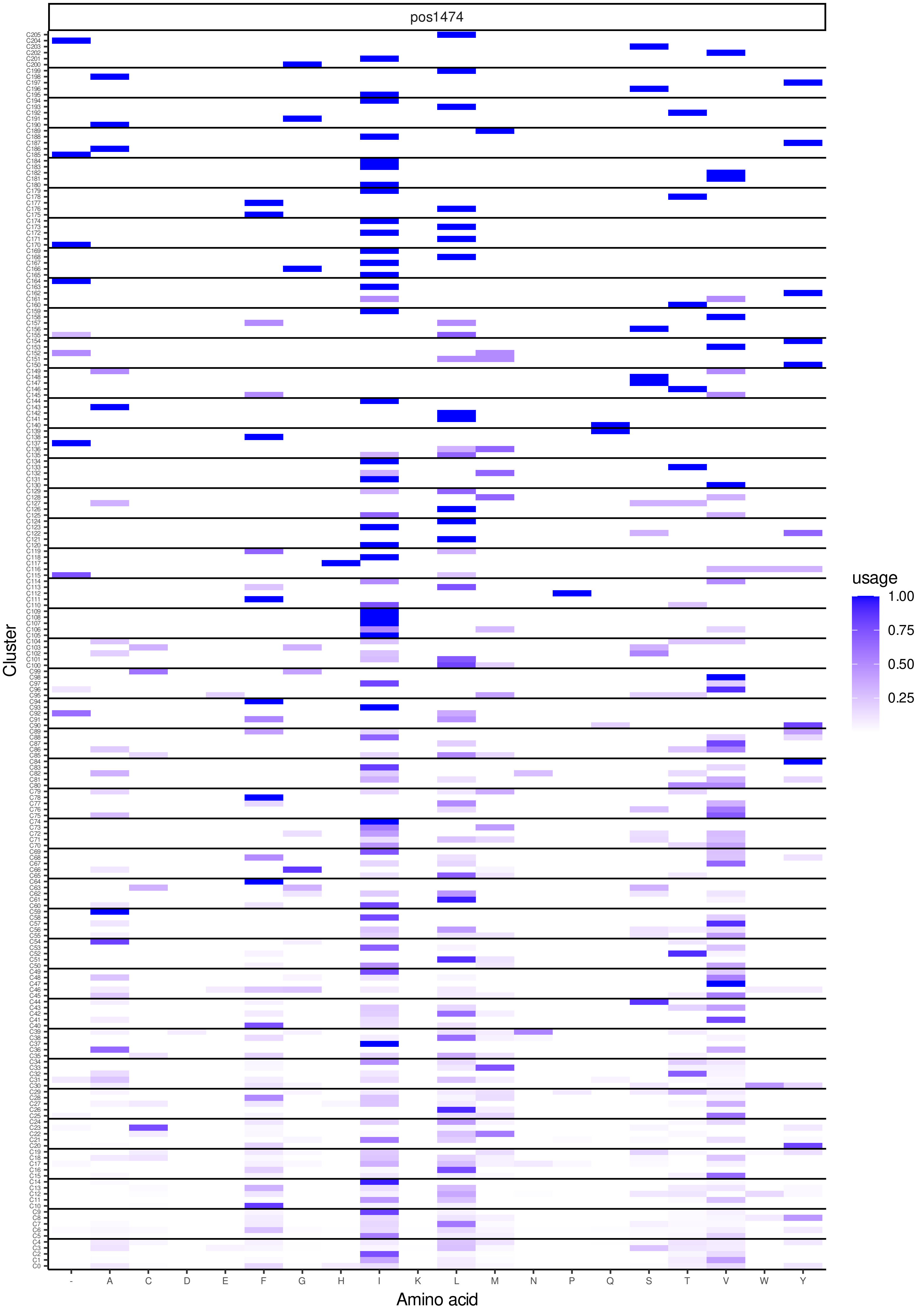

pos1474
Cluster
Amino acid
usage
1.00
0.75
0.50
0.25
- A C D E F G H I K L M N P Q S T V W Y

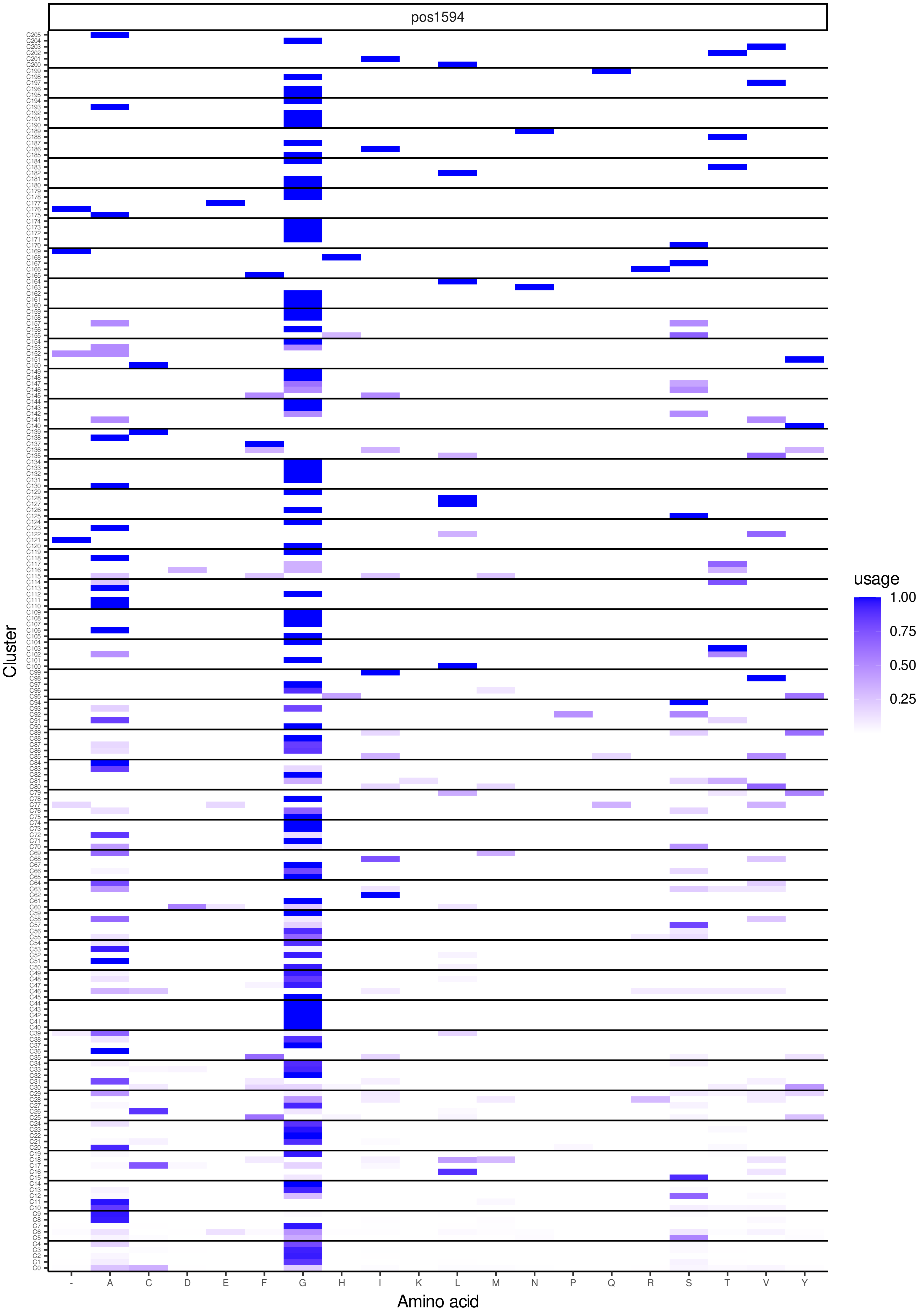
pos1594
Cluster
Amino acid
usage
1.00
0.75
0.50
0.25
-
A
C
D
E
F
G
H
I
K
L
M
N
P
Q
R
S
T
V
Y

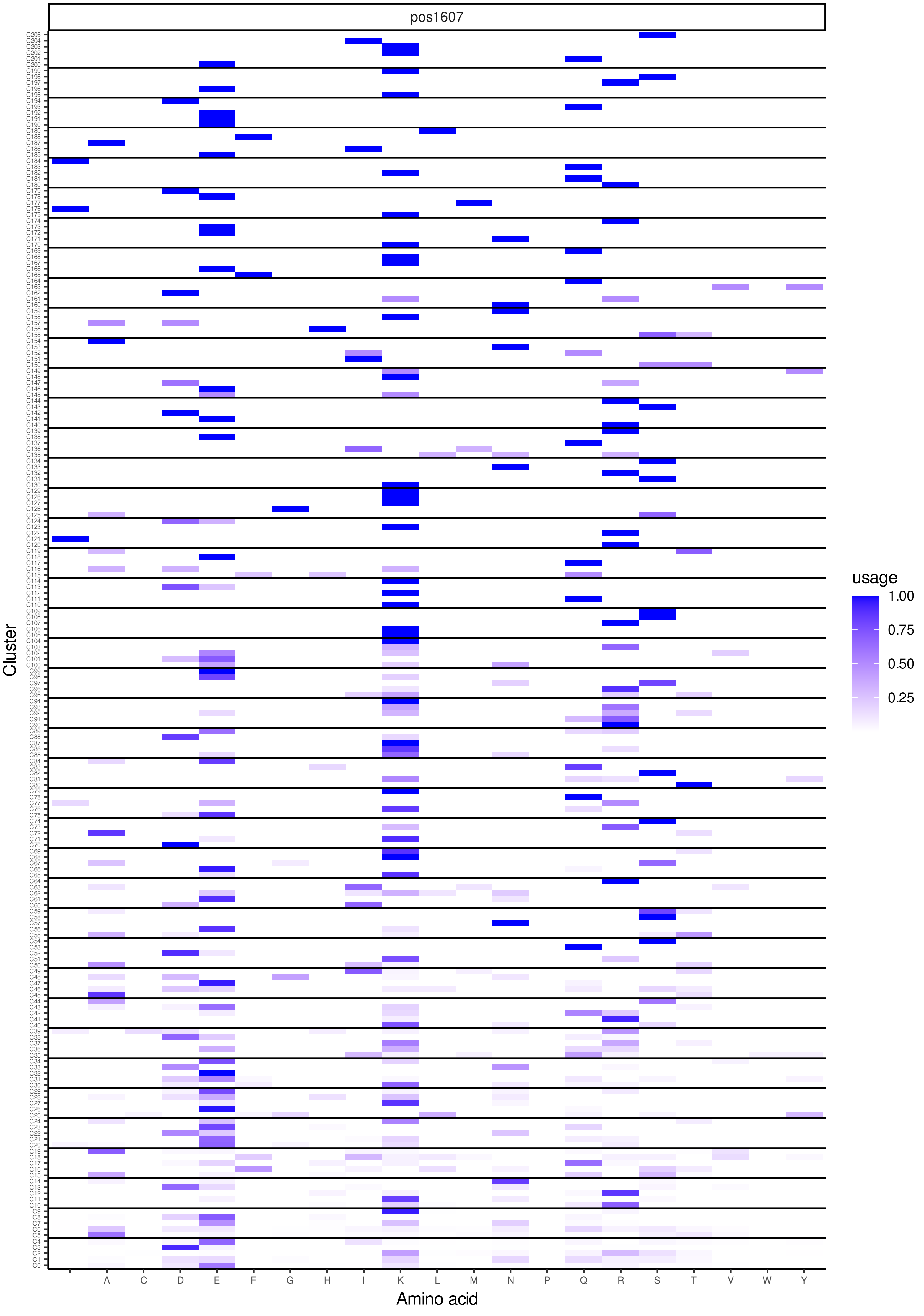
pos1607
Cluster
Amino acid
usage
1.00
0.75
0.50
0.25
- A C D E F G H I K L M N P Q R S T V W Y

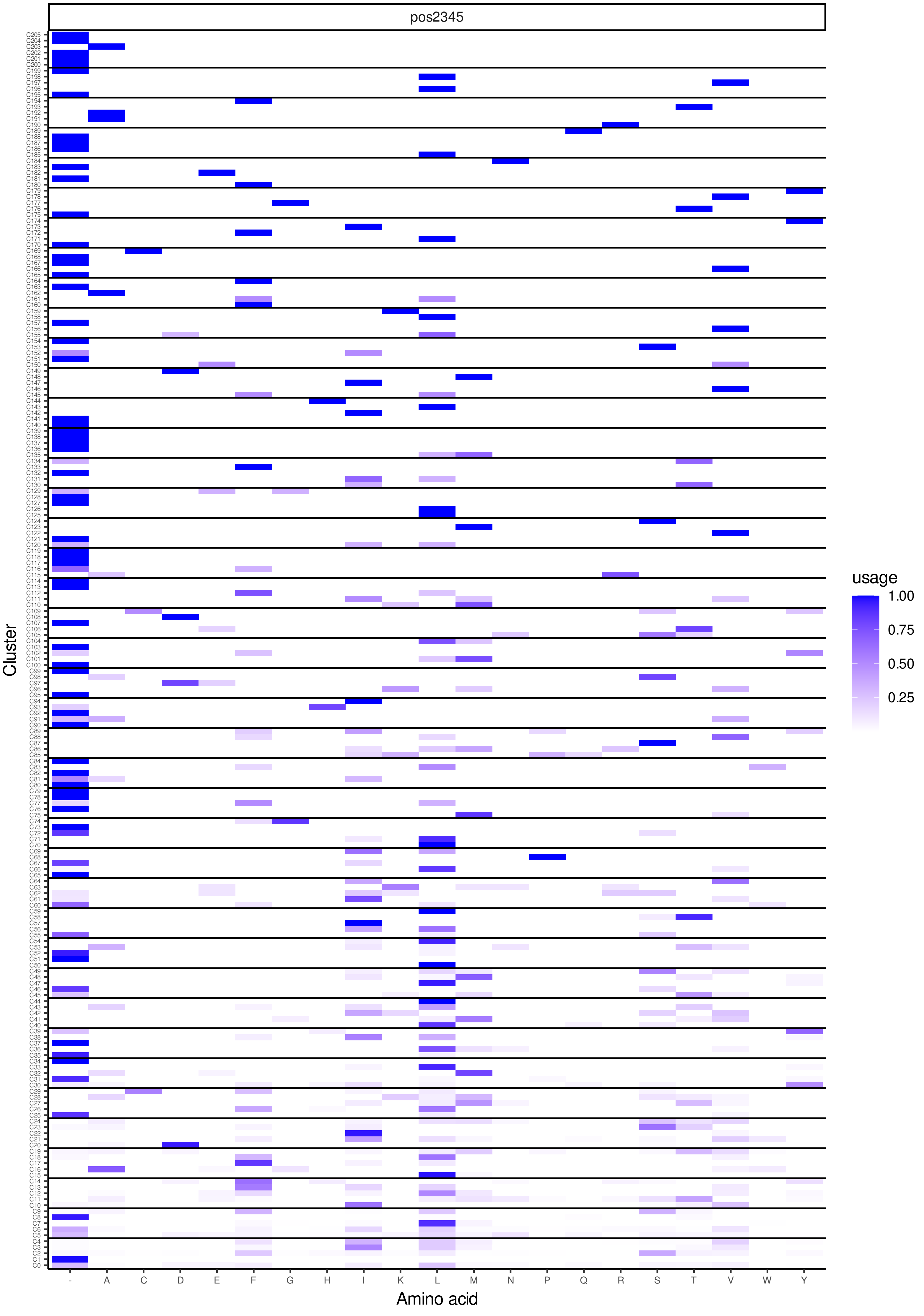

pos2345
Cluster
Amino acid
- A C D E F G H I K L M N P Q R S T V W Y
usage
1.00
0.75
0.50
0.25

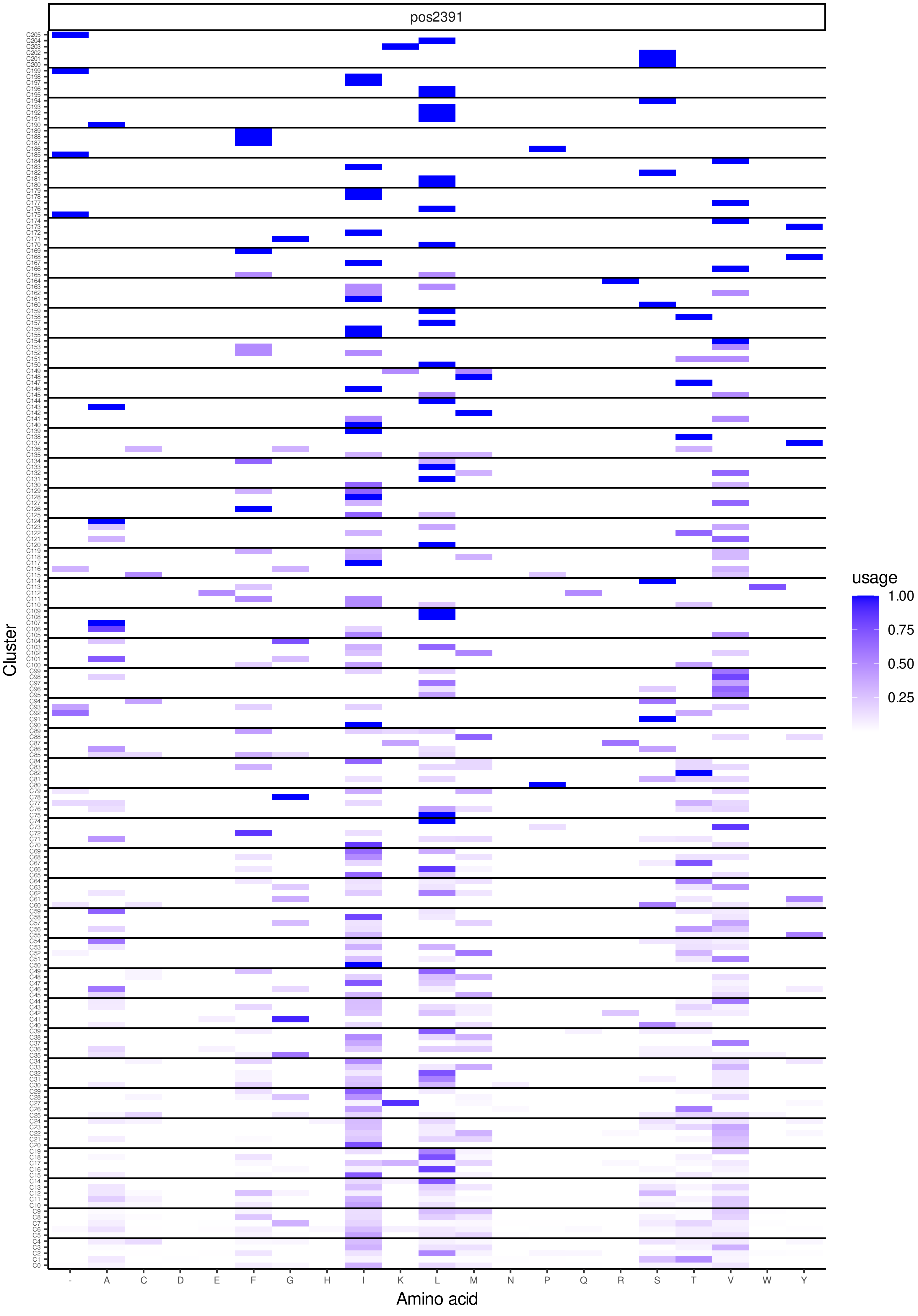
pos2391
Cluster
Amino acid
- A C D E F G H I K L M N P Q R S T V W Y
usage
1.00
0.75
0.50
0.25
C0 … C205

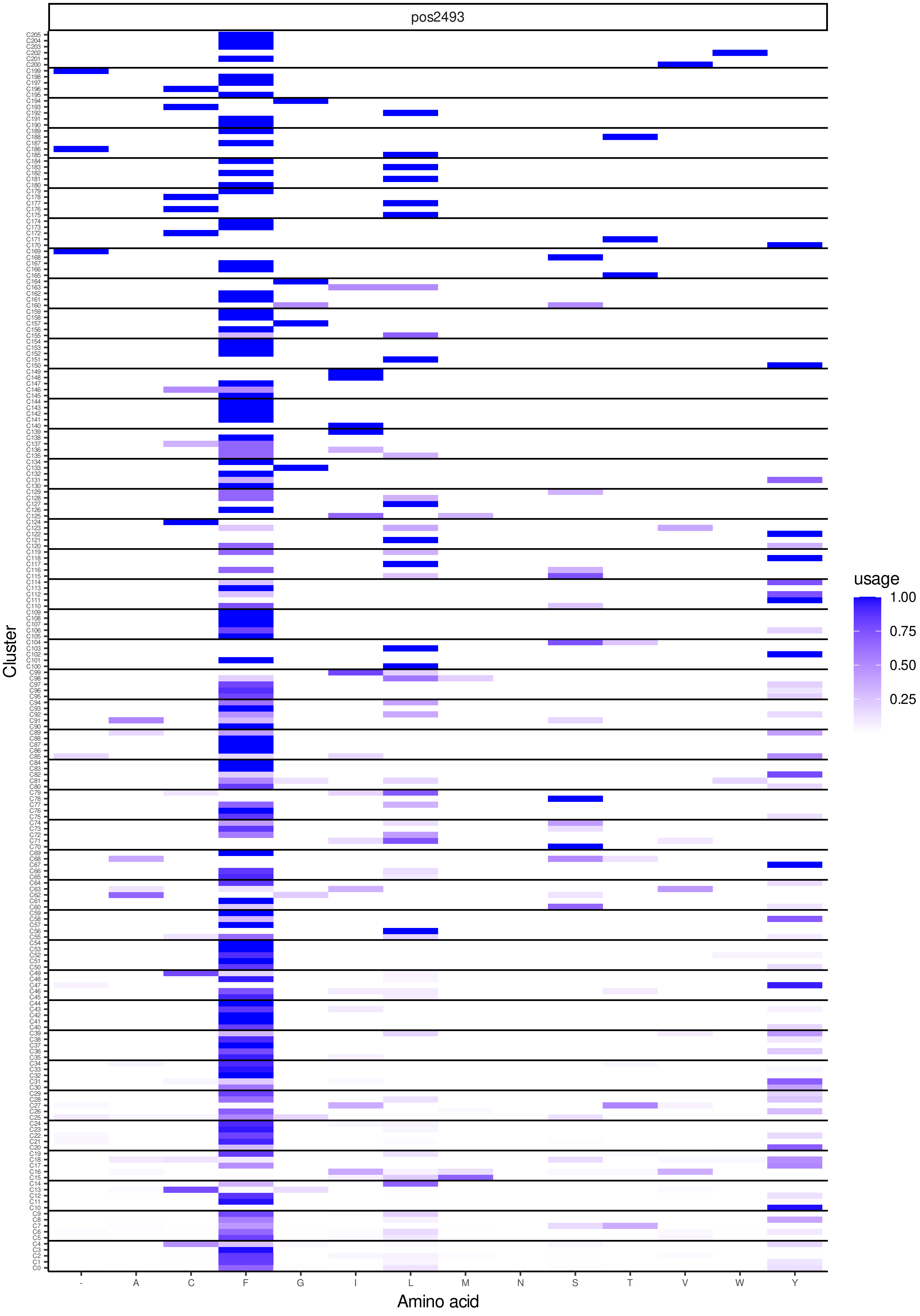
pos2493
Cluster
Amino acid
- A C F G I L M N S T V W Y
usage
1.00
0.75
0.50
0.25

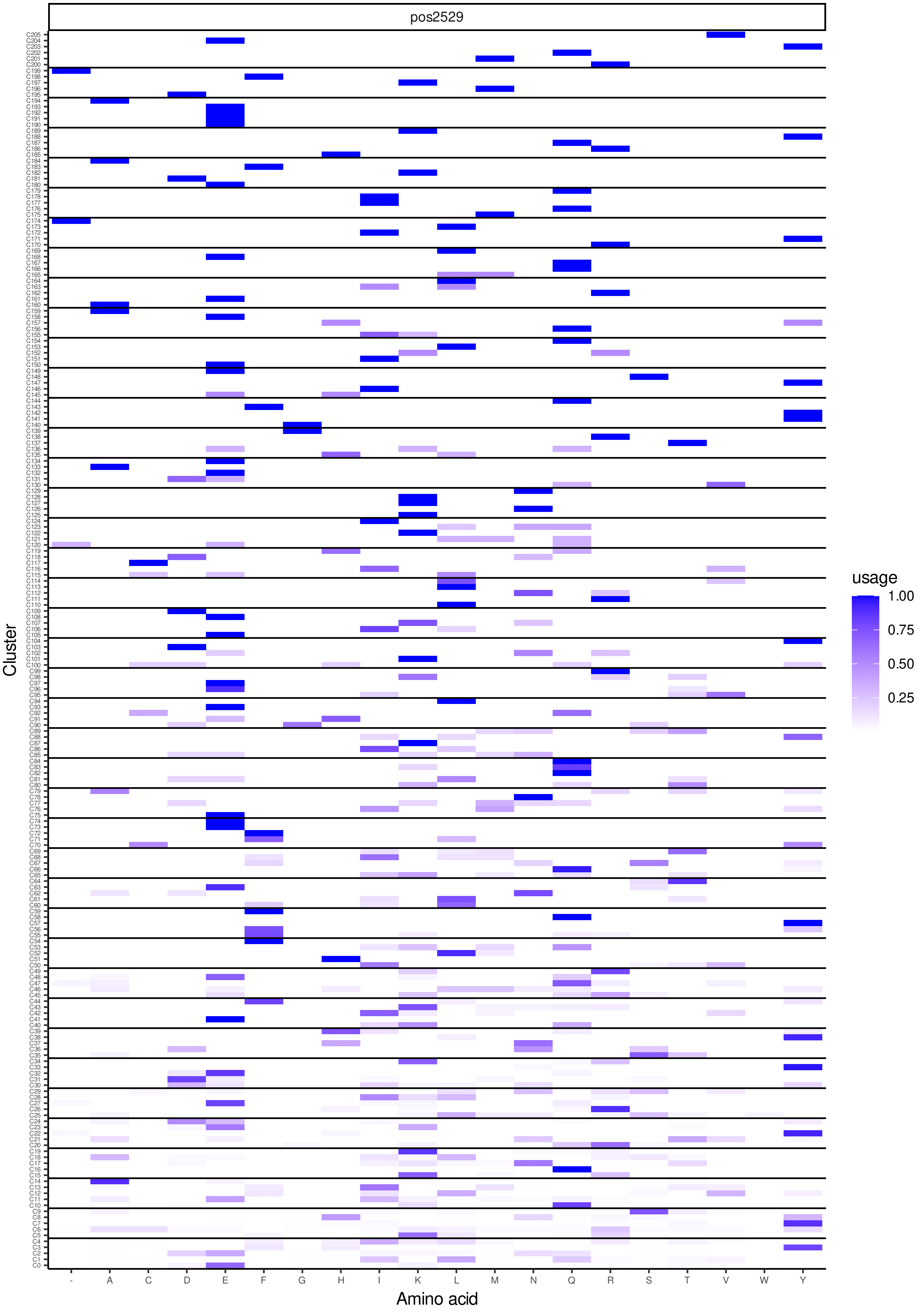

pos2529
Cluster
Amino acid
- A C D E F G H I K L M N Q R S T V W Y
usage
1.00
0.75
0.50
0.25

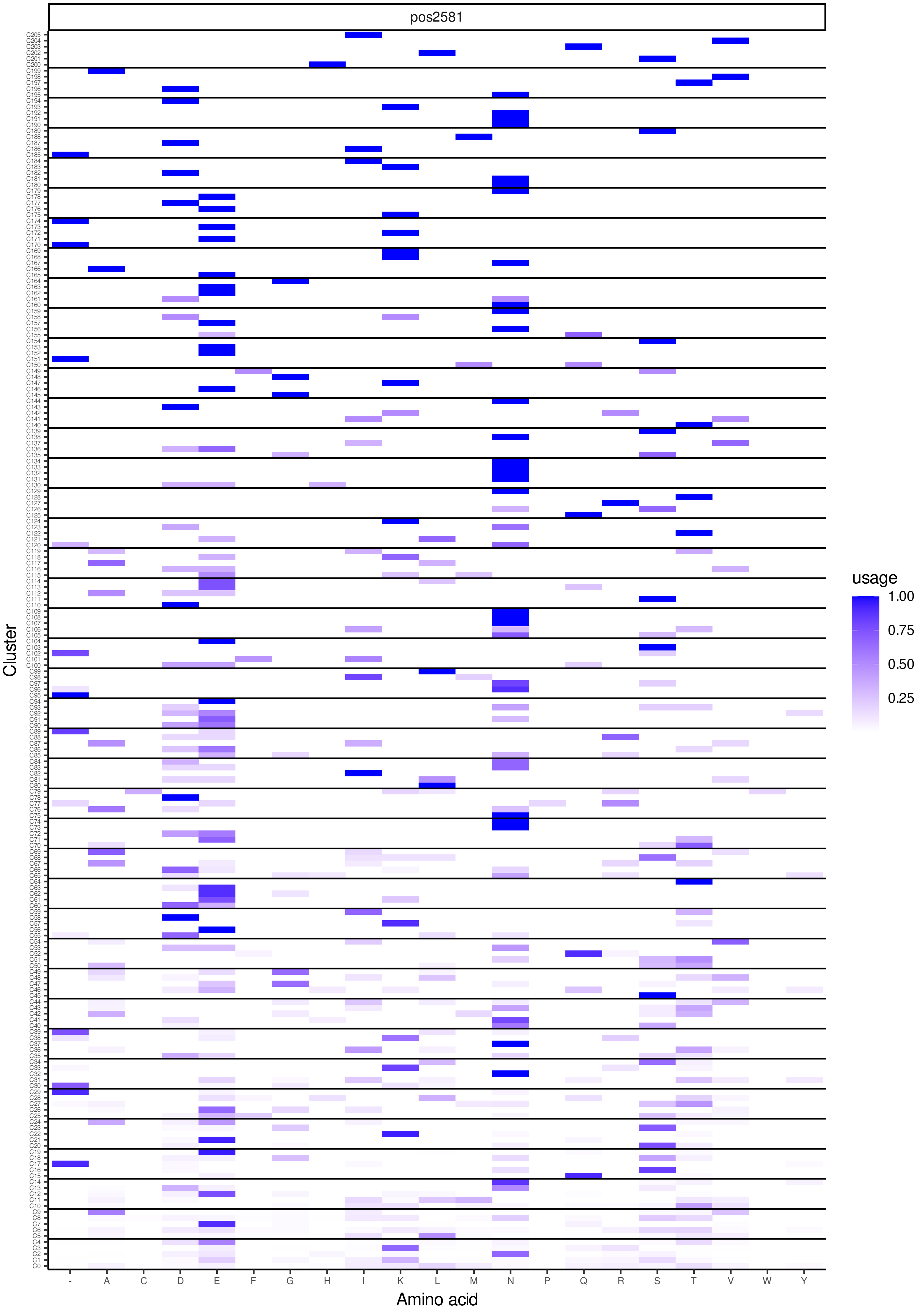
pos2581
Cluster
Amino acid
usage
1.00
0.75
0.50
0.25
- A C D E F G H I K L M N P Q R S T V W Y

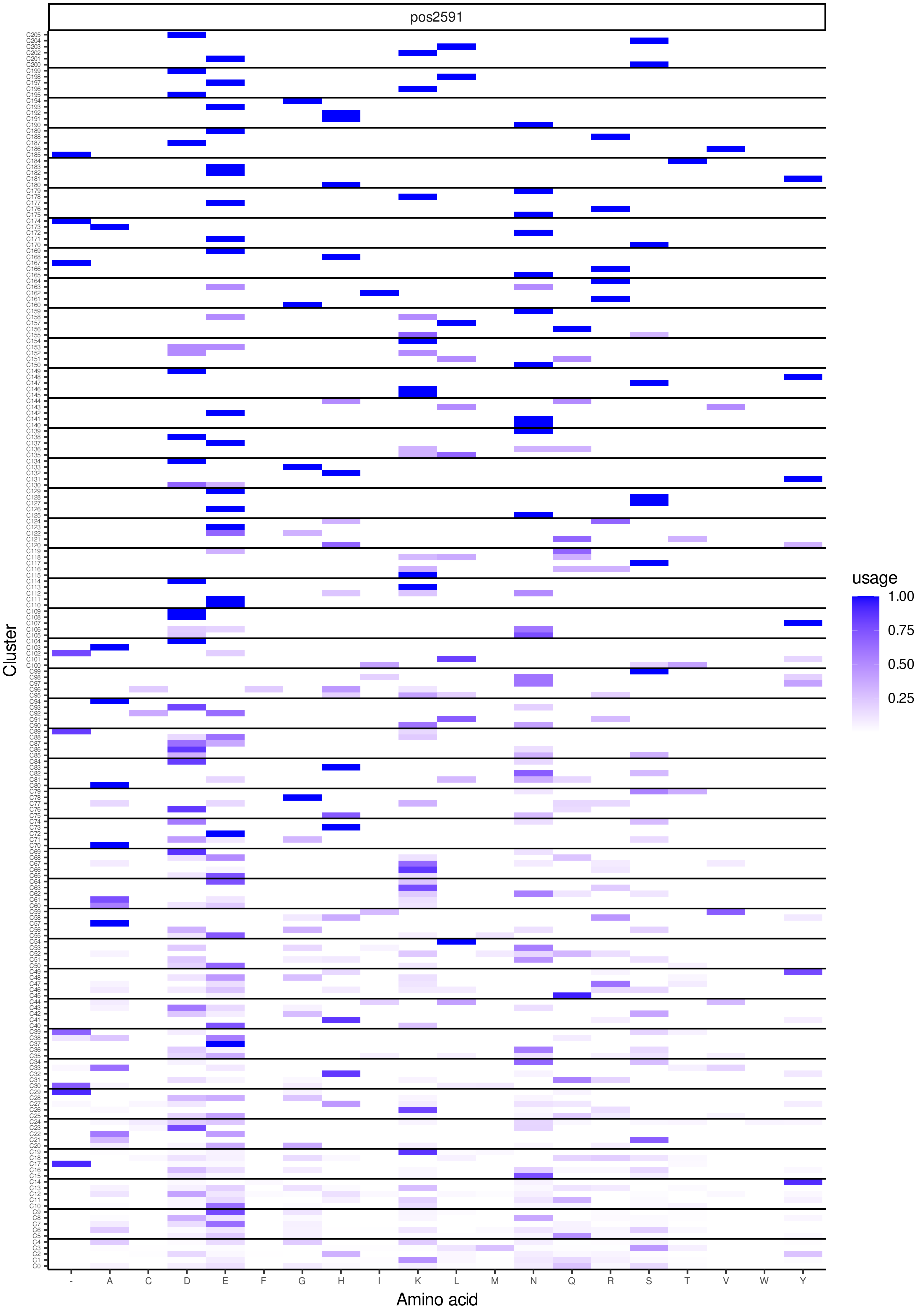
pos2591
Cluster
Amino acid
- A C D E F G H I K L M N Q R S T V W Y
C0 … C205
usage
1.00
0.75
0.50
0.25

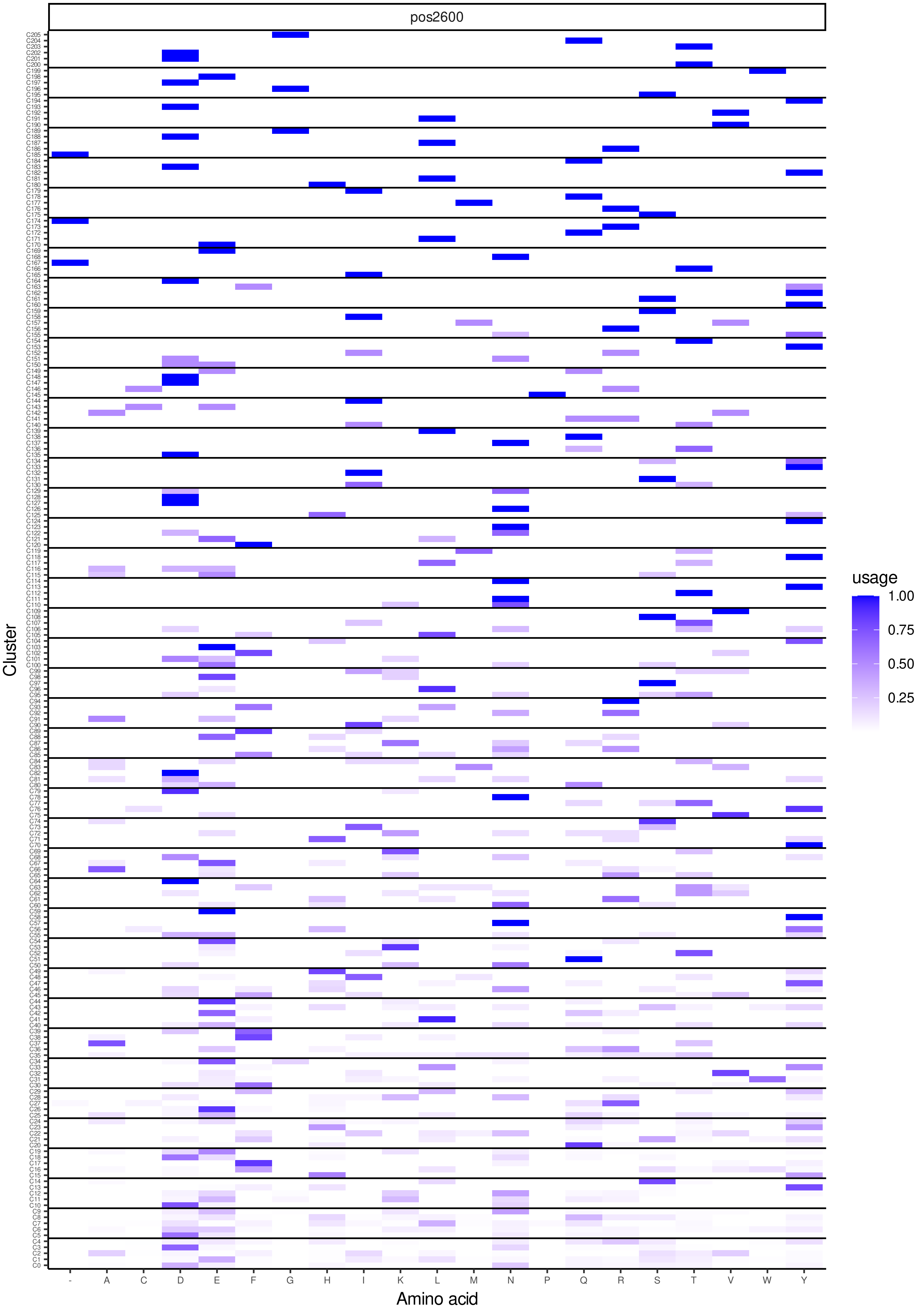
pos2600
Cluster
Amino acid
- A C D E F G H I K L M N P Q R S T V W Y
usage
1.00
0.75
0.50
0.25

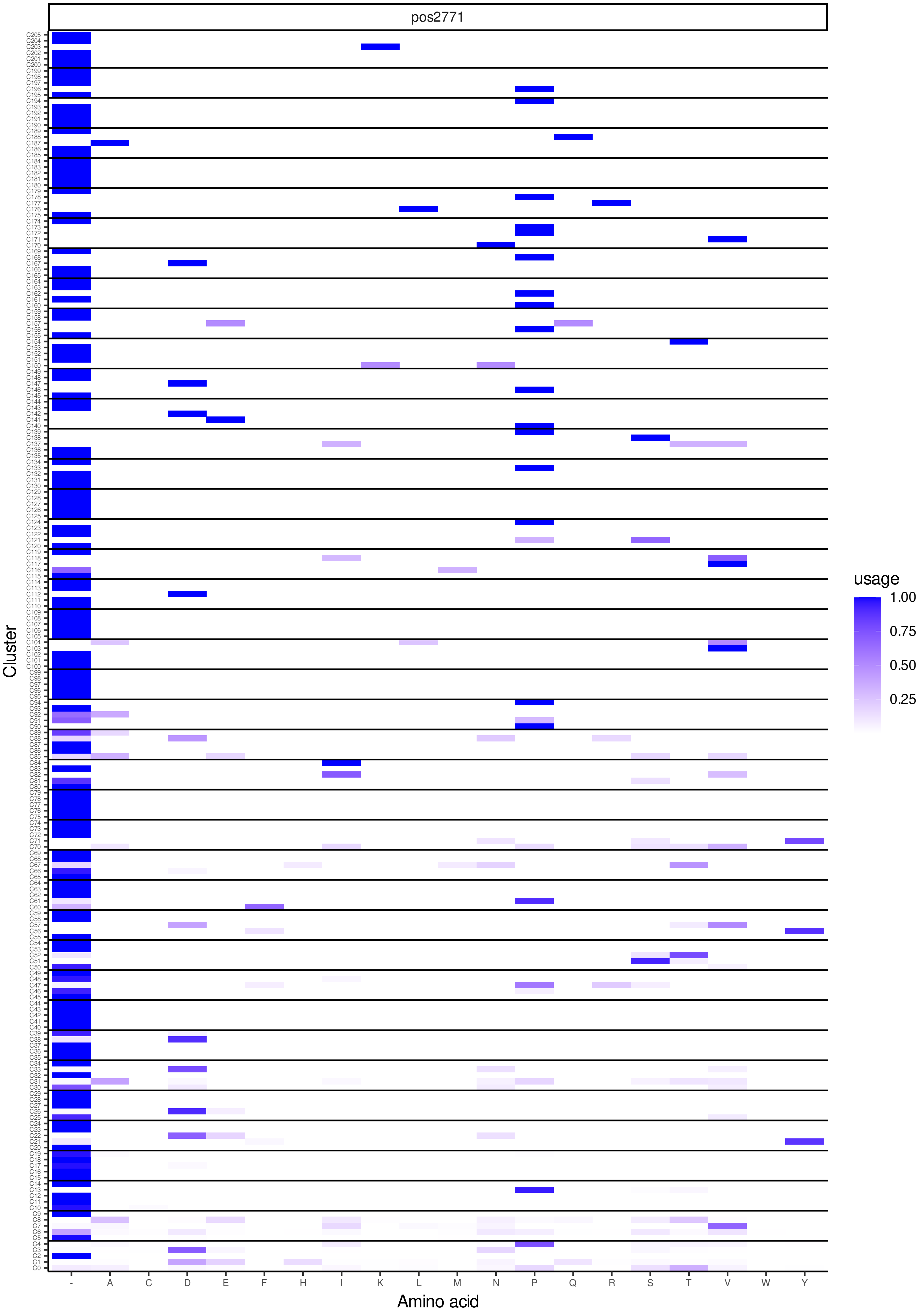
pos2771
Cluster
Amino acid
usage
1.00
0.75
0.50
0.25
- A C D E F H I K L M N P Q R S T V W Y

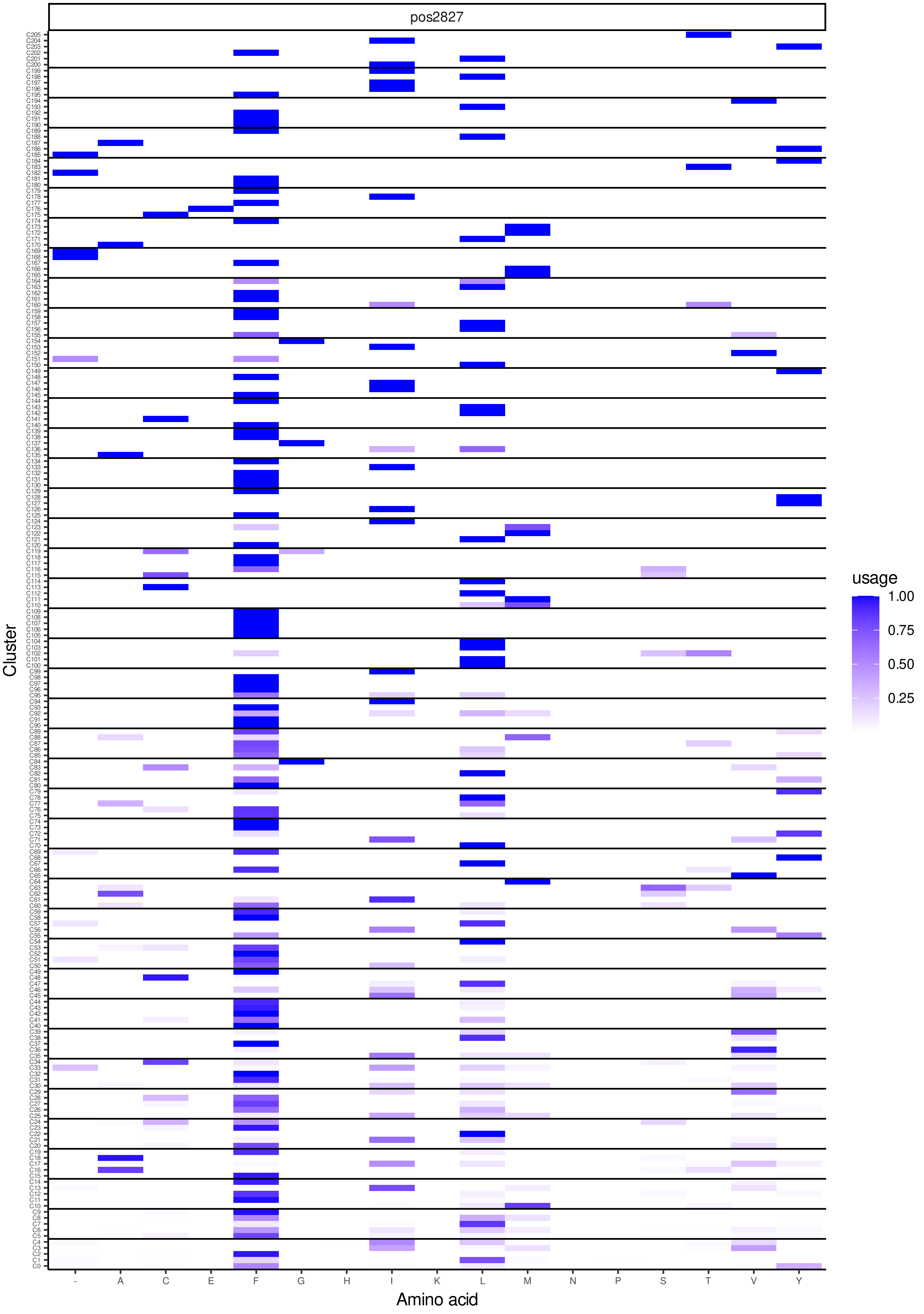
pos2827
Cluster
Amino acid
- A C E F G H I K L M N P S T V Y
usage
1.00
0.75
0.50
0.25
